# Human inhalable antibody fragments neutralizing SARS-CoV-2 variants for COVID-19 therapy

**DOI:** 10.1101/2021.06.04.447066

**Authors:** Olga Minenkova, Daniela Santapaola, Ferdinando Maria Milazzo, Anna Maria Anastasi, Gianfranco Battistuzzi, Caterina Chiapparino, Antonio Rosi, Giuseppe Gritti, Gianmaria Borleri, Alessandro Rambaldi, Clélia Dental, Cécile Viollet, Bruno Pagano, Laura Salvini, Emanuele Marra, Laura Luberto, Antonio Rossi, Anna Riccio, Emilio Merlo Pich, Maria Gabriella Santoro, Rita De Santis

## Abstract

As of October 2021, coronavirus disease 2019 (COVID-19) caused by the severe acute respiratory syndrome coronavirus 2 (SARS-CoV-2) remains a global emergency, and novel therapeutics are urgently needed. Here we describe human single chain variable fragment (scFv) antibodies (76clAbs) that block an epitope of the SARS-CoV-2 spike protein essential for ACE2-mediated entry into cells. 76clAbs neutralize the delta variant and other variants being monitored (VBMs) and inhibit spike-mediated pulmonary cell-cell fusion, a critical feature of COVID-19 pathology. In two independent animal models, intranasal administration counteracted the infection. Due to high efficiency, remarkable stability, resilience to nebulization and low production cost, 76clAbs may become a relevant tool for rapid, self-administrable early intervention in SARS-CoV-2-infected subjects independently of their immune status.

## Main Text

After almost two years since the SARS-CoV-2 pandemic began, although vaccination is starting to show its positive effect, the medical community is still urgently calling for novel cost-effective therapeutic and prophylactic tools to control infection in not-immunized as well as in vaccinated subjects partially protected against emerging variants (*1-3*). SARS-CoV-2 entry into the cells is mediated by docking of the spike, a trimeric class I fusion protein anchored into the viral envelope, on the human angiotensin converting enzyme 2 (hACE2) (*4,5*) and several spike-blocking anti-SARS-CoV-2 monoclonal antibodies (mAbs) are being sought for passive immunization (*6*). Three mAbs received Emergency Use Authorization (EUA) from the Food and Drug Administration (FDA) for the treatment of mild to moderate COVID-19 in non-hospitalized patients and dozens are in clinical trials. Nevertheless, current evidence is still insufficient to draw meaningful conclusions regarding the utility of injected SARS-CoV-2- neutralizing mAbs (*7*) whose wide spread use is hampered by at least three limitations: first, loss of neutralization capacity towards emerging variants (*8*); second, health system’s sustainability because of the costs of these products administered at gram doses and produced in expensive mammalian expression systems; third, the risk of Antibody Dependent Enhancement of the infection (ADE) (*9*). Additionally, recent data indicate that, after the systemic infusion of a neutralizing mAb, SARS-CoV-2 is still present in the nasal turbinates (*10*), where the virus initially harbors and from where it spreads, making intranasal and aerosol treatments particularly attractive (*11*). Small-size, single domain V_H_H nanobodies, specific for SARS-CoV-2, were proposed for a topical use i.e. by inhalation, as an alternative to systemic mAbs (*12,13*). However, being nanobodies of camelid origin, they require sophisticated humanization procedures to avoid immunogenic responses, potentially hampering their full development (*14*). Here we present novel small-size human antibodies in the format of single-chain variable fragments (scFv) that share several advantages with nanobodies with fewer risks. The scFv antibodies were generated by engineering variable heavy and light chain sequences of immunoglobulins in a single polypeptide by joining them via a flexible peptide linker that allows reconstitution of the functional antigen binding domains (**Fig. 1A)** (*15*). Being devoid of the immunoglobulin Fc, scFv antibodies are not at risk of inducing ADE and, due to their rapid production at large scale in non-mammalian expression systems, they could be developed at relatively lower cost compared to mAbs. Moreover, because of their smaller size (about 28 kDa vs the 150 kDa of mAbs) and low complexity (no glycosylation, simpler structure), they are particularly stable and suitable for topical use in non-hospital settings. For example, the first human scFv antibody, brolucizumab, was recently approved for topical treatment of exudative age-related macular degeneration by FDA, thus paving the regulatory pathway of this class of molecules (*16*). In this article we describe SARS-CoV-2-neutralizing human scFv antibodies that were obtained by panning phage display libraries on the spike receptor binding domain (RBD). For maximizing the possibility to select affinity-matured antibody species, we profiled the immune response to SARS-CoV-2 of ten COVID-19 convalescent donors from the Bergamo hospital, Italy. As shown in **Fig. 1B, left panel**, six out of ten COVID-19 sera (CS) were found to strongly bind to the SARS-COV-2 spike (Wuhan strain). These six sera also proved to be the most potent at inhibiting the spike-hACE2 interaction (**Fig. 1B, right panel**) and the best to neutralize viral infectivity in Vero E6 cells (table S1). Total RNA, extracted from lymphocytes of the six best responders, was converted to cDNA and the immunoglobulin genes were amplified and cloned in scFv antibody format for phage display, as depicted in fig. S1A. After selection cycles by panning, binders were subjected to affinity maturation by error-prone mutagenesis of the heavy chain and to light chain shuffling followed by repeated panning on RBD under stressing conditions (low antigen concentration, low pH washing, high temperature). Selected scFv5, scFv86 and scFv76 antibodies, as well as affinity matured derivatives from scFv76 (i.e., scFv76-46, scFv76-55, scFv76-57, scFv76-58 and scFv76-77), here coded as scFv76-cluster antibodies (76clAbs), all binding spike in the low nanomolar range, were produced in soluble form and purified according to the procedure schematized in fig. S1B. The amino acid sequences of CDR3 and germline gene repertoire of the selected scFv antibodies are reported in table S2 that shows 76clAbs having identical VH CDR3 originating from the IGVH3-53 germline, with minor differences in the remaining VH sequence. Diversity is contributed by VL sequences that are derived from the IGVK3-20 germline except for scFv76-55 and scFv76-57 antibodies that originate VL from IGVK3-15 and IGVK1-9 germlines, respectively. Moreover, scFv5 and scFv86 antibodies share identical VH, derived from the IGVH1-46 germline, and have VL derived from the IGVL2-14 and IGVK3-11 germlines, respectively. Recombinant spike S1-subunit and RBD-domain spike proteins of the SARS-CoV-2 variant of concern (VOC) delta and of variants being monitored (VBMs), all associated with high infectivity and immune escape, were then used to test scFv antibody reactivity in terms of binding, affinity and inhibition of spike/hACE2 interaction. We included in the analyses spikes carrying the following mutations: D614G, here indicated as S1D614G, present now in all viral isolates; T19R, G142D, E156G, 157-158d, L452R, T478K, D614G, P681R of the delta variant (S1delta); HV69-70del, Y144del, N501Y, A570D, D614G, P681H of the alpha variant (S1alpha); K417N, E484K, N501Y and D614G, of the beta variant (S1beta). RBD proteins with single or double mutations of most frequent variants were also tested. Results, shown in fig. S2 and table S3, collectively indicate that 76clAbs strongly bind to mutated proteins and recognize much more efficiently all spikes and RBD mutated proteins as compared to scFv5 and scFv86 antibodies that, based on the pattern of reactivity, appear to recognize a different antigenic epitope. In a set of independent experiments, binding affinity of scFv antibodies for SARS-CoV-2 spike was measured by Surface Plasmon Resonance (SPR). Kinetic data on mutated spike and RBD proteins are reported in **Table 1**, fig. S3, S4 and table S4, showing *K*_*D*_ values ranging between sub-nanomolar and 1-digit nanomolar concentrations, except for scFv76-55 that exhibits a *K*_*D*_ of 14.9 nM for S1beta. SPR analyses did not reveal additive binding of the 76clAbs, suggesting that they might recognize the same or a very close antigenic epitope, while scFv5 binds to an independent epitope (fig. S5A). To test the ability of 76clAbs to recognize a SARS-CoV-2 native spike, HEK293T cells were transfected with the WT S1 spike-encoding plasmid. High content screening (HCS) fluorescent imaging showed scFv76 antibody binding to HEK293T spike+ cells (fig. S5B). Similar data were obtained with all 76clAbs and confirmed by cytofluorimetry indicating the ability of such antibodies to recognize the spike expressed on the cell surface (fig.S5C). The 76clAbs were then evaluated for their ability to inhibit the binding of the viral spike to hACE2 receptor and for their capacity to neutralize viral infectivity. Results in **Fig. 1C** show that 76clAbs, but not scFv5 or scFv86, blocked spike/hACE2 interaction and that, except for scFv76-55, they were highly resilient to mutations of the delta variant, as well as to other mutations of VBMs (fig. S6). The IC_50_ values of 76clAbs, ranging from 0.36 to 4.30 nM, are reported in table S5. Interestingly, and consistently with binding and SPR affinity data, these results indicate that 76clAbs, except for scFv76-55, substantially maintain the capacity to inhibit the binding of the spike variants and mutated RBD to hACE2.

**Fig. 1.**
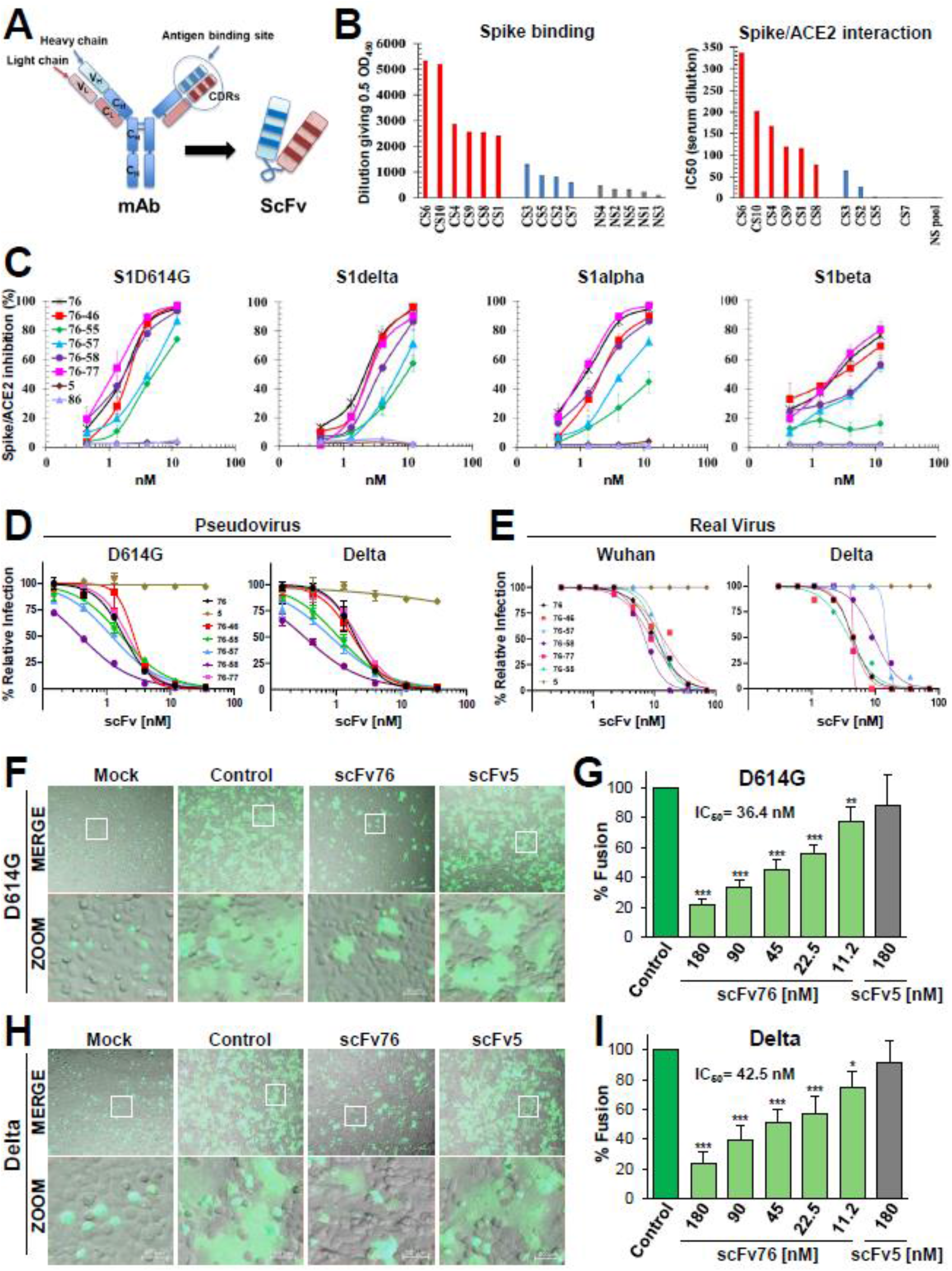
Selection and functional characterization of anti-SARS-CoV-2 human scFv antibodies. (**A**) Schematic representation of scFv and mAb. (**B**) Analysis of COVID-19 convalescent (CS) and negative control (NS) sera for binding to SARS-CoV-2 spike (left) and inhibition of spike/hACE2 interaction (right) by ELISA. Data expressed as the dilution giving 0.5 OD and as IC_50_, respectively. (**C**) Inhibition of interaction of indicated spikes with hACE2 measured by ELISA. Data are the average (±SE) of 2-3 experiments. (**D**) Neutralization of pseudotyped-virus expressing indicated spikes assessed by luciferase-assay in hACE2-expressing Caco-2 cells. Data are the average (±SD) of two replicates from one representative experiment. (**E**) Microneutralization assay of SARS-CoV-2 Wuhan and delta strains in Vero E6 cells. (**F-I**) Inhibition of SARS-CoV-2 spike-mediated cell-cell fusion using HEK293T donor cells expressing GFP and D614G-mutated (**F**,**G**) or Delta (**H**,**I**) spike, or GFP only (mock), incubated 1 h with saturating dose (180 nM) of scFv76 or scFv5 (**F**,**H**) or with scalar doses of scFv76 (**G**,**I**), and then overlaid on monolayers of hACE2-expressing A549 cells for 4 hrs. Scale bar 200 μm, zoom 50 μm. Cell-cell fusion quantification expressed as percentage relative to control (average ±SD of 5 fields from two biological replicates). *p<0.05, **p<0.01, ***p<0.001; ANOVA test

**Table 1.** Affinity of anti-SARS-CoV-2 scFv antibodies for spike protein variants by Surface Plasmon Resonance.

| scFv | Spike S1 variant | $k_a$<br>($10^5 \text{ M}^{-1} \text{ s}^{-1}$) | $k_d$<br>($10^{-5} \text{ s}^{-1}$) | $K_D$<br>(nM) |
| --- | --- | --- | --- | --- |
| **76** | WT | 1.4 | 4.7 | 0.3 |
|  | S1D614G | 1.3 | 6.0 | 0.4 |
|  | S1alpha | 1.1 | 4.3 | 0.4 |
|  | S1beta | 1.3 | 54.0 | 4.1 |
|  | S1delta | 1.0 | 6.6 | 0.7 |
| **76-46** | WT | 1.9 | 3.1 | 0.2 |
|  | S1D614G | 1.9 | 4.8 | 0.3 |
|  | S1alpha | 1.4 | 7.3 | 0.5 |
|  | S1beta | 1.1 | 83.7 | 7.7 |
|  | S1delta | 1.0 | 4.6 | 0.4 |
| **76-55** | WT | 1.4 | 42.1 | 3.0 |
|  | S1D614G | 1.5 | 39.9 | 2.7 |
|  | S1alpha | 1.2 | 61.6 | 5.1 |
|  | S1beta | 1.5 | 223.0 | 14.9 |
|  | S1delta | 1.0 | 54.2 | 5.6 |
| **76-57** | WT | 1.5 | 7.6 | 0.5 |
|  | S1D614G | 1.5 | 6.7 | 0.4 |
|  | S1alpha | 1.1 | 10.6 | 0.9 |
|  | S1beta | 1.5 | 49.5 | 3.3 |
|  | S1delta | 0.9 | 9.4 | 1.1 |
| **76-58** | WT | 2.8 | 32.6 | 1.1 |
|  | S1D614G | 3.3 | 34.7 | 1.1 |
|  | S1alpha | 2.8 | 34.0 | 1.2 |
|  | S1beta | 1.7 | 75.0 | 4.5 |
|  | S1delta | 1.8 | 37.6 | 2.1 |
| **76-77** | WT | 1.5 | 3.0 | 0.2 |
|  | S1D614G | 1.4 | 4.9 | 0.3 |
|  | S1alpha | 1.4 | 2.0 | 0.1 |
|  | S1beta | 1.1 | 31.6 | 2.8 |
|  | S1delta | 0.9 | 4.7 | 0.5 |
| **5** | WT | 0.5 | 19.8 | 4.3 |
|  | S1D614G | 0.4 | 17.5 | 4.4 |
|  | S1alpha | 0.5 | 19.8 | 4.2 |
|  | S1beta | 0.5 | 20.0 | 4.1 |
|  | S1delta | 0.3 | 15.8 | 4.9 |
| **86** | WT | 0.7 | 9.6 | 1.5 |
|  | S1D614G | 0.6 | 11.4 | 1.8 |
|  | S1alpha | 0.7 | 11.2 | 1.7 |
|  | S1beta | 0.7 | 11.5 | 1.6 |
|  | S1delta | 0.5 | 10.4 | 2.1 |

To investigate the potential escape of virus variants to 76clAbs neutralization, human colon Caco-2 cells stably expressing the hACE2 receptor (Caco-2 hACE2) were infected with luciferase-expressing SARS-CoV-2 S-pseudoviruses bearing the D614G mutation or the delta spike variant. Results in **Fig. 1D** confirmed that 76clAbs, but not scFv5, inhibited the infection of both pseudoviruses and the infection of pseudoviruses bearing spike mutations of several VBMs (fig. S7A). Consistently with previous neutralization data, scFv76-55 was the least effective of the cluster and the most sensitive to mutations. Similar results were obtained comparing the neutralization potency of scFV76 and scFV76-55 on pseudoviruses bearing the alpha, beta and gamma spike variants (fig. S7B). Neutralization of infectivity of the real SARS -CoV-2 virus was then tested by microneutralization assay of cytopathic effect (CPE) in Vero E6 cells. Interestingly, all 76clAbs proved to be effective with MN50 (50% microneutralization titer) <15 nM against the delta variant (**Fig. 1E**). Neutralization potency of scFv76, scFv76-46 and scFv76-58 antibodies was then further evaluated by RT-qPCR in human pulmonary Calu-3 cells infected with the SARS-CoV-2 Wuhan strain, D614G, or alpha variants. Results in table S6 show that when the 76clAbs were added to the cell culture 1 h after the infection they were effective at inhibiting all virus strains with IC_50_ below 22 nM, while when added 1 h before infection, IC_50_ was below 2.2 nM. Similar difference in the neutralization potency before or after infection was observed with the immune sera pooled from the six COVID-19 convalescent donors of **Fig. 1B** and table S1.

The ability of 76clAbs to prevent SARS-CoV-2 spike-induced fusion of pulmonary cells was tested *in vitro*. This analysis has high translational value, since syncytia formation in the lung is considered a most relevant pathologic hallmark of COVID-19, driving localized inflammation and thromboembolism (*17*). Incubation with nanomolar concentrations of the scFv76 antibody proved to be strongly effective at inhibiting fusion between WT (Wuhan) (fig. S8), D614G (**Fig. 1F and G**) and delta (**Fig. 1H and I**) spike-expressing human HEK293T cells and human lung A549 cells stably expressing the hACE2 receptor (A549 hACE2). These observations were replicated in monkey Vero E6 target cells (fig. S8).

Identification of the antigenic epitope recognized by anti-SARS-CoV-2 scFv antibodies was performed by high-throughput flow cytometry on human HEK-293T cells expressing Wuhan or alanine mutated RBD proteins (alanine scanning library of 194 aa). Data in **Fig. 2A** show for scFv76, scFv76-55, scFv76-57, scFv76-58 and scFv86, cytofluorimetry dot plots indicating primary (red) and secondary (blue) critical clones bearing alanine substitutions in positions strongly reducing binding. Such critical residues (red and blue spheres) are visualized on a crystal of the SARS-CoV-2 spike protein trimer (PDB ID # 6XCN) and the SARS-CoV-2 spike protein receptor binding domain (PDB ID# 6Z2M). Results indicate that 76clAbs recognize an epitope on the spike RBM that includes F456, Y473, N487 and Y489 residues, while the critical residue for the non-neutralizing scFv86 is mapped outside RBM. Interestingly, none of the residues composing the epitope recognized by 76clAbs has been, so far, found mutated neither in the delta VOC nor in VBMs (fig. S9). From alanine scanning data it is possible to speculate that such epitope might be invariant because, being structurally located at the tip of the spike, its conformation is essential for the docking to hACE2. This epitope was found to be restricted to SARS-CoV-2 as indicated by lack of reactivity of 76clAbs with SARS-CoV, MERS-CoV and HCoV-HKU1 spikes (**Fig. 2B**). Overall, the data so far collected regarding functional profiles of 76clAbs together with epitope mapping collectively show that the reactivity of these antibodies is not affected by mutations present in SARS-CoV-2 variants, including those previously shown to be associated to the escape of antibody recognition (*8, 18, 19*).

**Fig. 2.**
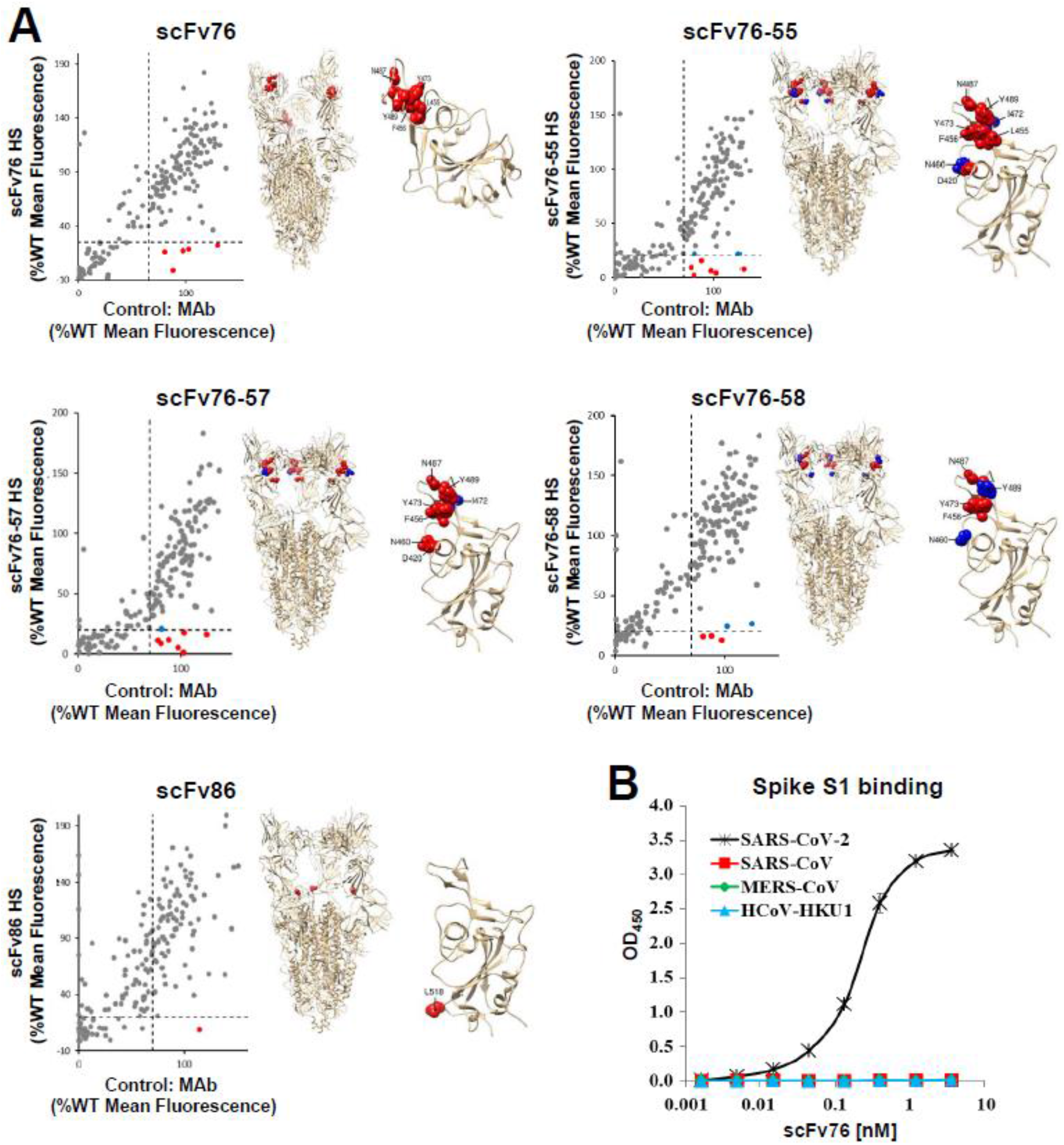
Epitope mapping of anti-SARS-CoV-2 human scFv antibodies. **(A)** Flow cytometry dot plot representation of antibody binding to alanine-mutated clones (left). For each point, background fluorescence was subtracted from the raw data, which were then normalized to reactivity with WT target protein. To identify preliminary primary critical clones (red circles), a threshold (dashed lines) of >65% WT binding to control Ab and <25% WT binding to test Abs was applied. Secondary clones (blue circles) are highlighted for clones that did not meet the set thresholds but whose decreased binding activity and proximity to critical residues suggested that the mutated residue may be part of the antibody epitope. Critical residues (red spheres) for scFv antibodies binding, and secondary residues (blue spheres) that may contribute to binding, are also visualized on crystal structure of the SARS-CoV-2 spike protein trimer (PDB ID # 6XCN) (center) and on SARS-CoV-2 spike protein receptor binding domain (PDB ID # 6Z2M) (right). (**B**) Reactivity of scFv76 towards indicated spikes by ELISA.

Regarding the biochemical properties of 76clAbs, size exclusion chromatography (SEC-HPLC) and SDS-PAGE electrophoresis (**Fig. 3A** and fig. S10) indicated an apparent molecular weight of about 28 kDa, as expected based on deduced amino acid sequence. Molecular weight was further confirmed by SEC-HPLC-mass spectrometry (MS) using 20 mM ammonium formate as native elution buffer. The average spectrum obtained from all MS spectra of the scFv76 antibody is reported in **Fig. 3B**, where two gaussian dispersions, centered on m/z 2800 and m/z 4000, are present. Their deconvolution indicates the presence of two species with molecular weight of 28008 and 56016 that correlate with the scFv76 monomeric and dimeric forms, respectively. The result of peptide mapping performed by trypsin digestion of scFv76, scFv76-46, scFv76-58 and scFv76-77, followed by reverse phase-ultra-HPLC-MS/MS analysis, confirmed 100% matching of peptides with the expected amino acidic sequence. Thermal resistance, selected as a surrogate marker of protein stability, was then investigated. The 76clAbs were subjected to heating for 1 h at increasing temperature and tested for antigen binding by ELISA. Results in **Fig. 3C** identify scFv76, scFv76-46 and scFv76-58 as the most resistant antibodies, and scFv76-57 as the most sensitive one. To assess protein folding and secondary structure content, far-UV circular dichroism (CD) spectra of 76-clAbs were recorded at 20 and 90 °C (**Fig. 3D**). CD spectra of antibodies at 20 °C resemble those of folded proteins with significant β-sheet content, while the spectra at 90 °C show that the antibodies have undergone unfolding. Quantitative analysis of CD spectra at 20 °C, by means of BeStSel (*20*), indicates that 76clAbs have a folded structure with about 40% of antiparallel β-sheet content (table S7). Considering the possible use of the 76clAbs for aerosol therapy of COVID-19, their resilience to nebulization stress was finally evaluated. Data showed that aqueous solutions of scFv76, at milligram/mL concentrations, can be nebulized by mesh nebulizer, maintaining unaltered both protein integrity (**Fig. 3E**) and spike binding property (**Fig. 3F**).

**Fig. 3.**
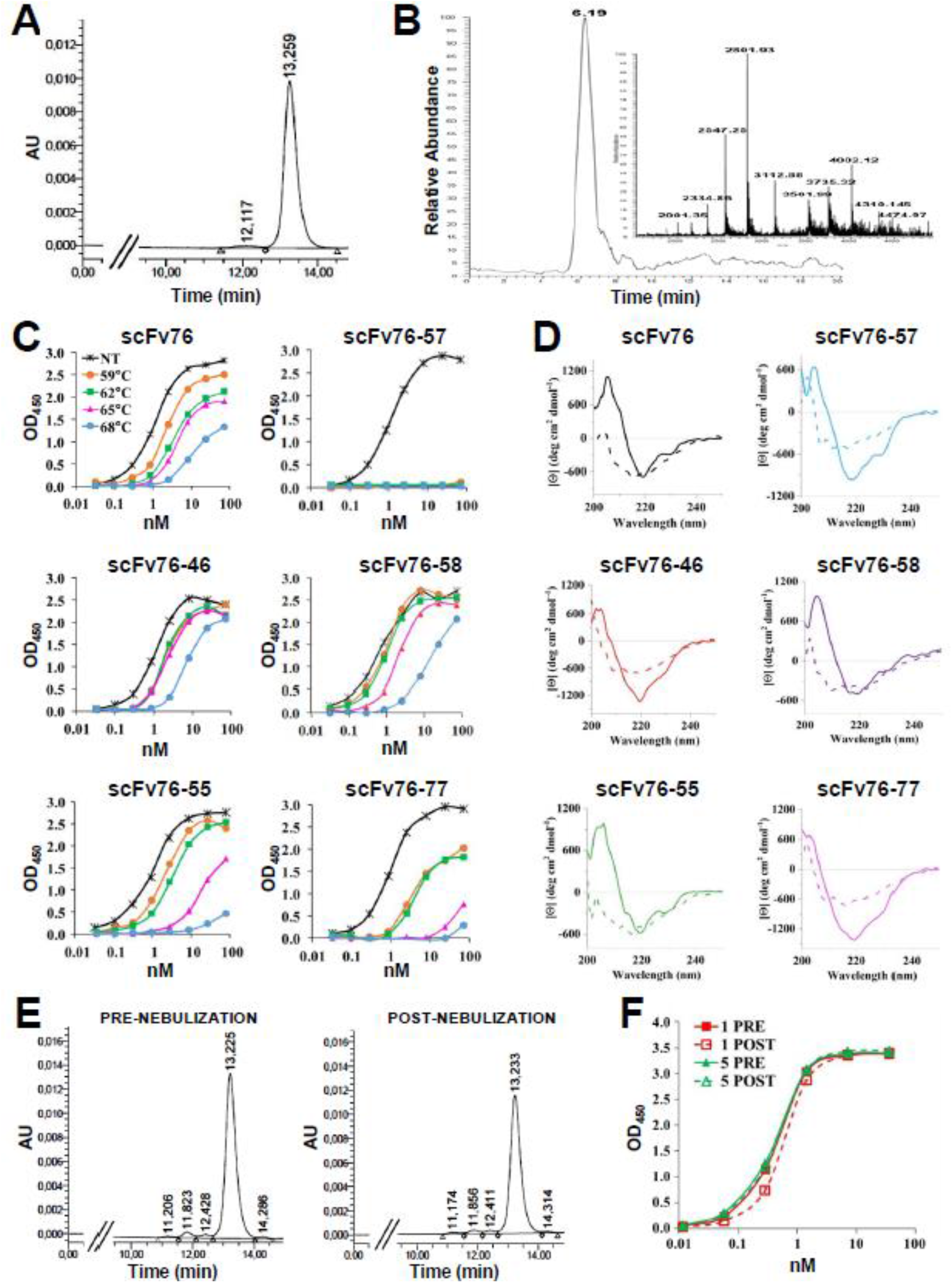
Biochemical and structural characterization of 76clAbs. (**A**) UV chromatogram at 280 nm from SEC-HPLC analysis of scFv76 in 50 mM phosphate buffer, 150 mM NaCl, 10% acetonitrile, pH 7.2 on TSKgel G3000 SWXL 30 cm x 7.8 mm column (Tosoh Bioscience). (**B**) Total ionic current (TIC) chromatogram from SEC-HPLC analysis of scFv76 in 20 mM ammonium formate, pH 6.8 on Mab Pac SEC-1 column (Thermo). Inset, mass spectrum as average of spectra recorded between 5 and 7 min. (**C**) Binding of thermally stressed and not treated (NT) scFv76-clAbs. (**D**) Far-UV circular dichroism spectra of antibodies. Analysis was performed in PBS buffer at 20 (solid lines) and 90 (dashed lines) °C. (**E**) UV chromatograms at 280 nm from SEC-HPLC analysis, as in **A**, of scFv76 pre- and post-nebulization at 1 mg/ml. (**F**) Binding of scFv76, pre- and post-nebulization at 5 mg/mL or 1 mg/mL, to SARS-CoV-2 RBD by ELISA. Data are the average (± SD) of three replicates from one representative experiment.

In vivo effects of the topical administration of various members of 76clAbs against SARS-CoV-2 infection was then investigated in a model of hACE2-expressing transgenic (tg) mice, infected with a luciferase-expressing D614G S-pseudotyped virus. Infection levels in nasal turbinates were measured by bioluminescence imaging (BLI). As shown in **Fig. 4A**, the intranasal administration of 74 μg/mouse of scFv76 (37 μg/nostril), 2 h before and 4 h after infection, completely blocked the viral infection as deduced by the loss of luminescent signal (**Fig. 4B)**. In a confirmatory study, scFv76-58 and one unrelated scFv antibody were also tested. Results, shown in **Fig. 4C**, confirmed efficacy of both 76clAbs, while the unrelated antibody was not active, thus indicating treatment specificity. In an independent study, hamsters, one of the few animals sensitive to human SARS-CoV-2, were infected with the D614G SARS-CoV-2 strain. Hamsters were intranasally administered with 70 μg of scFv76 (35 μg/nostril), 2 h before and once daily for the two consecutive days post-infection. Results showed protection from body weight loss (**Fig. 4D**) and reduction of nasal discharge (**Fig. 4E**). Finally, propaedeutic investigation of scFv76 aerosol delivery was performed by nose-only exposure of mice (fig. S11A) to antibody solutions nebulized by the Aerogen Pro (Aerogen) mesh nebulizer, a nebulization device used for human treatment. Results in fig. S11B indicated that nebulized scFv76 antibody can be effectively delivered to the lung down to the alveolar space, as shown by immunohistochemistry. The amount of antibody deposited appears to increase with time of exposure, as shown by tissue staining intensity (**Fig. 4F**) and by RBD binding ELISA of homogenized lungs (**Fig. 4G)**.

**Fig. 4.**
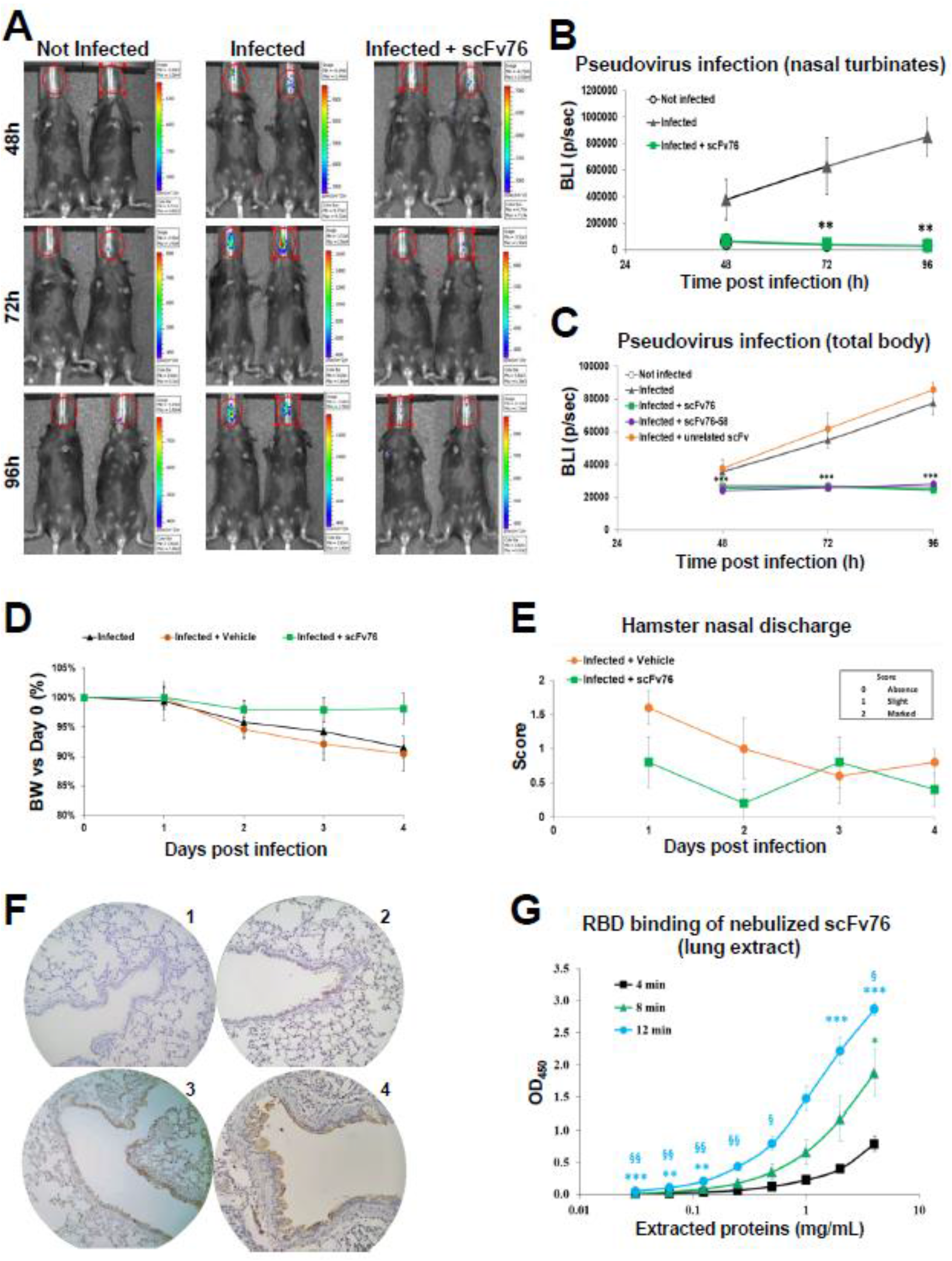
ScFv76 antibody inhibits viral infectivity in animal models by intranasal administration. **(A**) Representative bioluminescence images (BLI) after intranasal administration of scFv76 in k18-hACE2 mice infected with a luciferase-expressing SARS-CoV-2 spike (D614G) pseudotyped-virus. Imaging by Xenogen IVIS200. (**B**) Light photons/sec in nasal turbinates and (**C**) total body. BLI data expressed as mean (± SE) of 5 animals. ***p< 0.001 and **p< 0.01; Student’s t-test versus vehicle-treated mice. (**D**) Body weight and (**E**) nasal discharge of hamsters intranasally infected with SARS-CoV-2 (D614G) 10^5^ pfu/animal on day 0 (D0) and treated with ScFv76 or vehicle 2 hours before and once daily for 2 days post-infection. Body weight expressed as % versus D0. Nasal discharge scored as in the inset. Data are mean (±SD) of 5 animals/group. (**F**) Immunohistochemistry of pulmonary sections of mice, 1h after PBS (panel 1) or 4-(panel 2), 8-(panel 3) or 12-min (panel 4) scFv76 (5 mg/mL solution) aerosol exposure. (**G**) SARS-CoV-2 RBD binding by ELISA of lung proteins extracted from mice as in **F**. Data expressed as mean (± SD) (n=4). ***p< 0.001, **p< 0.01 and *p< 0.05; Student’s t-test versus 4-min-nebulized mice. ^§§^p< 0.01 and ^§^p< 0.05; Student’s t-test versus 8-min-nebulized mice.

Because of reported SARS-CoV-2 molecular mimicry of human proteins and related extensive cross-reactivity of anti-virus antibodies with human normal tissues (*21*), scFv76 was tested by immunohistochemistry on tissue microarrays (fig. S12A). Representative pictures in fig. S12B clearly show reactivity of an anti-PDL1 scFv (generated by the same technology as scFv76 and equally detected by anti-flag or anti-his tag antibodies) with several tissues while no reactivity was observed with scFv76, thus confirming its suitability as a potential drug for human use. In summary, here we describe a novel class of anti-SARS-CoV-2 human scFv antibodies endowed with subnanomolar neutralization potency, in vivo efficacy and significant stability. These small size fully human antibodies are expected to be well tolerated and be produced at low costs thus being promising candidates for early treatment of COVID-19. Remarkably, the ability of 76clAbs to recognize an essential and conserved spike antigenic epitope translates into the neutralization of the delta variant and the so far identified VBMs, some of which known to drive escape of approved and investigational mAbs (*22*). The single chain antibodies here proposed for nasal or lung topical delivery might overcome limitations of parenteral mAbs aimed for a systemic passive immunization (*23*). Developed as a complementary approach to the existing vaccination and passive systemic immunization, that do not induce a robust mucosal defense (*24*), this class of antibodies may contribute to halt the SARS-CoV-2 infection at its early stages.

## Supporting information

Suppl Mat

## Supplementary Materials

Materials and Methods

Tables S1 to S7

Figs. S1 to S12

References (25-35)

## Acknowledgments

We are grateful to COVID-19 convalescent subjects for providing blood samples. We thank Integral Molecular, Philadelphia, USA for expedite identification of the antibody epitope; Dr. Silvia Santopolo for help with pseudovirus RNA titration; Dr Nestor Santiago Gonzalvo (Cytiva) for excellent technical assistance on Biacore analyses; Mr Claudio Albertoni and Mrs Evelyn Vaccaro for expert technical assistance in animal models. Prof Luigi Giusto Spagnoli for expert opinion on immunohistochemistry.

## Author contributions

Conceptualization: GG, GB^2^, AR^2,3^, MGS^8,9^, EMP, OM and RDS; Investigation: DS, AMA, FM, BP, GB^1^, LS, CC, AR^1^, CD, CV, AR^8^, LL, AR^9^; Supervision and manuscript writing: MGS^8,9^, EMP and RDS.

## Data and materials availability

Antibodies described in the paper can be provided upon material transfer agreement (MTA) subscription.

