## Supplementary material for "Human inhalable antibody fragments neutralizing SARS-CoV-2 variants for COVID-19 therapy": Suppl Mat

*ScFv antibodies against a spike conserved epitope and neutralizing virus variants are suitable for nose and lung protection*

Olga Minenkova[1], Daniela Santapaola[1], Ferdinando Maria Milazzo[1], Anna Maria Anastasi[1], Gianfranco Battistuzzi[1], Caterina Chiapparino[1], Antonio Rosi[1], Giuseppe Gritti[2], Gianmaria Borleri[2], Alessandro Rambaldi[2,3], Clélia Dental[4], Cécile Viollet[4], Bruno Pagano[5], Laura Salvini[6], Emanuele Marra[7], Laura Luberto[7], Antonio Rossi[8], Anna Riccio[9], Emilio Merlo Pich[1], Maria Gabriella Santoro[8,9] and Rita De Santis[1*]

[1]Alfasigma SpA, Biotechnology R&D, Pomezia, Rome, Italy; [2]Papa Giovanni XXIII Hospital, Bergamo, Italy; [3]Department of Hematology and Oncology, University of Milan, Italy; [4]Texcell EVRY Cedex, France; [5]Department of Pharmacy, University of Naples Federico II, Italy; [6]Toscana Life Sciences; [7]Takis Biotech, Rome, Italy; [8]Institute of Translational Pharmacology, CNR, Rome, Italy; [9]Department of Biology, University of Rome Tor Vergata, Rome, Italy

*Corresponding author
Alfasigma SpA, Biotechnology R&D Via Pontina Km 30.400 Pomezia 00071, Rome, Italy

**Supplementary Materials**
Materials and Methods
Tables S1 to S7
Fig. S1 to S12
References (25-35)

### Materials and Methods

#### Convalescent sera and mRNA

Total RNA was purified from 10 mL of peripheral blood of ten COVID-19 convalescent subjects, confirmed virus-free by double negative swab, among the personnel of ASST Papa Giovanni XXIII Hospital of Bergamo. Matched sera from each donor were also provided. All samples were anonymized, and donors consented to the use of their biological materials for the generation of phage libraries displaying antibodies on their surface, according to a protocol approved by the Ethical Committee.

#### Phage libraries

Total RNA was converted to cDNA by using Super Script IV VILO Master Mix (Thermo Fisher Scientific). For library construction cDNAs derived from lymphocytes of the best 6 responders, CV1, CV4, CV6, CV8, CV9 and CV10, were used. Gene fragments encoding for immunoglobulin variable domain were amplified with specific primers and assembled into scFv antibody genes as described earlier (*25*). The libraries of scFv format were prepared by inserting assembled scFv genes in the proprietary vector pKM19 (*26, 27*). Each library had an average size of about $5x10^7$ independent clones. Maturation libraries were generated on the base of originally selected clones by error-prone PCR amplification of heavy chains and by light chain shuffling. Heavy chain DNA fragment of the chosen clones was amplified with upstream vector primer and 3'-end VH primer. To introduce a low rate of mutations in the heavy chains under error-prone amplification the following conditions were used: 1 mM dTTP, 1 mM dGTP, 1 mM dCTP, 0.2 mM dATP, 0.1 mM dITP, 7 mM $MgCl_2$ (including $MgCl_2$ from Taq buffer) and 0.5 mM $MnCl_2$. The mutated VH DNA sequences were assembled with sequences encoding whole repertoire of VL by introducing between the two chains a short linker of 15 amino acids $(GGGGS)_3$ (*10*) and scFv encoding sequences were cloned in pKM19.

**ScFv selection**

About $5\times10^{11}$ TU (transducing units) of freshly amplified scFv antibody library were preincubated, in blocking buffer, with 10% of human serum, 10% of AD202 bacterial extract and 10% UV-killed M13KO7 for 30 min at 37 °C. Fifty µL of protein G conjugated Dynabeads were incubated with 2 µg of SARS-CoV-2 RBD human Fc for 30 min at room temperature (RT) with slow agitation, then washed, blocked with complete blocking buffer and added to the library and incubated for 1 h at 37 °C under gentle agitation. The bound phage was captured by using magnetic concentrator, washed ten times with PBS/Tween (PBS, 0.1% Tween20) and then eluted by 0.1 M glycine, pH 2.2, for 10 min at RT and then neutralized by 2 M Tris, pH 9.6. Conditions of selection of matured clones were designed to be more stringent. The phage suspension was incubated for 1 h at 63 °C before adding to the antigen coated carrier. Moreover, bound phages were intensively washed with washing buffer 10 times and additionally with 0.1 M glycine, pH 4.0, before elution. Once identified, the scFv genes were re-cloned in pKM16 (*26*) for production of soluble antibodies.

**Lab-scale production and purification of soluble scFv**

A plasmid DNA encoding for scFv antibody was used to transform E. coli DH5αF' cells. A single colony was inoculated in 50 mL of LB containing 100 µg/mL ampicillin and 2% glucose and incubated at 37 °C overnight (ON). Next day, 10 mL of bacterial suspension were inoculated in 1 L of LB containing 2% glucose and 100 µg/mL ampicillin. The cells were grown up to OD 0.8. Bacteria were then collected by centrifugation. The pellet was resuspended in 1 L of LB containing ampicillin and 20 mM $MgCl_2$ and incubated for further 4.5 h at 30 °C. Bacteria were collected, and the pellet was resuspended in 100 mM Tris-HCl, pH 7.4, 10 mM EDTA for periplasmic protein extraction and incubated ON at 30 °C with agitation. Cell debris was removed, and the supernatant was subjected to filtration by using Cogent µScale Tangential Flow Filtration System. Buffer was exchanged to 20 mM phosphate buffer, pH 7.4, supplemented with 25 mM imidazole, and volume was reduced to 50 mL. Histidine-tagged scFv was purified by using a HisTrap crude prepacked ready-to-used column operating with AKTA Avant liquid chromatography system (GE). The antibody was further purified from endotoxins by using Triton X-114. To purify solution from residual Triton X-114, Pierce Detergent Removal Spin Columns (Thermo Scientific) were used, according to manufacturer's instruction. Samples were then filter sterilized and stored in aliquots at -80 °C.

**Binding and spike/ACE2 or spike/RBD competition**

For binding experiments, Nunc maxisorp plates-96 wells were coated with 100 µL/well of SARS-CoV-2 spike S1 [SARS-CoV-2 spike glycoprotein S1 sheep Fc-tag, his-tagged spike S1 (D614G), spike S1(HV69-70del/Y144del/ N501Y, A570D, D614G, P681H), spike S1(K417N, E484K, N501Y, D614G), Spike S1 (T19R, G142D, E156G, 157-158 deletion, L452R, T478K, D614G,

P681R)] or his-tagged RBD proteins WT or with single or double mutation at K417, N439, L452, Y453, I472, A475, T478, V483, E484, N501, A520 (all from Sino Biological) in PBS, at final concentration 0.5 µg/mL, ON at +4 °C. Plates were blocked with blocking solution (1% BSA in PBS/Tween ) for 2 h at RT. After washing, dilutions of antibodies were added in a volume of 100 µL/well and incubated for 1 h at 37 °C. Plates were washed 4x with PBS/Tween and incubated for 1 h at RT with 100 µL/well of an anti-Flag HRP-conjugated antibody (Sigma-Aldrich) diluted 1:5000 in blocking buffer. After 4 washings with PBS/Tween, 100 µL/well TMB substrate were added and plates incubated at RT in the dark. Reaction was stopped by adding 50 µL of 2 N $H_2SO_4$. Absorbance was recorded at 450 nm by means of SUNRISE spectrophotometer (TECAN).
Similar experiments were performed for assessing the binding specificity of 76clAbs by using SARS-CoV Spike/S1, MERS-CoV Spike/S1 or HCoV-HKU1 spike/S1 his-tagged proteins (all from Sino Biological).
For competition experiments, Nunc maxisorp plates-96 wells were coated with 100 µL/well of spike S1 [SARS-CoV-2 spike glycoprotein S1 sheep Fc-Tag, his-tagged spike S1 (D614G), spike S1(HV69-70del/Y144del/ N501Y, A570D, D614G, P681H), spike S1(K417N, E484K, N501Y, D614G), Spike S1 (T19R, G142D, E156G, 157-158 deletion, L452R, T478K, D614G, P681R)] or his-tagged RBD proteins WT or with single or double mutation at K417, N439, L452, Y453, I472, A475, T478, V483, E484, N501, A520 (all from Sino Biological) in PBS, at final concentration 1.0-0.25 µg/mL, ON at +4 °C. Plates were blocked with 300 µL/well of blocking solution for 2 h at room temperature. After washing, dilutions of antibodies were added in a volume of 50 µL/well at double concentration and, after 30 minutes incubation at 37 °C, 1.0-0.25 µg/mL of human ACE2 protein mouse Fc-tag (Sino Biological) was added and incubated for 1 h at 37 °C. Plates were washed 4x with PBS/Tween and then incubated for 1 h at RT with 100 µL/well of an anti-mouse Fc conjugated to alkaline phosphatase (Sigma-Aldrich), diluted 1:1000 in blocking buffer. After 4 washings, 100 µL/well pNpp substrate were added and plates incubated at RT in the dark. Absorbance was recorded at 405 nm by means of SUNRISE spectrophotometer (TECAN).

**Surface Plasmon Resonance (SPR)**

Kinetic constants were determined by SPR experiments through a Biacore T200 instrument (Cytiva). Wuhan strain SARS-CoV-2 spike S1 mouse Fc-tagged protein (Sino Biological, 5 µg/mL in acetate buffer, pH 5), mutated SARS-CoV-2 spike S1 his-tagged proteins (Sino Biological, 5 µg/mL in acetate buffer, pH 5) and SARS-CoV-2 RBD his-tagged proteins (Wuhan strain and mutated ones, Sino Biological, 2.5 µg/mL in acetate buffer, pH 5) were immobilized at 350 RU level (S1 proteins) and at 100 RU level (RBD proteins) on the surface of a flow cell of a Series S sensor chip CM5 (Cytiva) by classical amine coupling procedure, while another flow cell surface was blank-immobilized with ethanolamine to be used as control surface. Then scFv proteins were flowed at 30 µL/min on both flow cells at 0.47 nM, 1.40 nM, 4.19 nM, 12.56 nM, 37.67 nM and 113 nM concentrations in HBS-EP+ buffer (Cytiva) for a contact time of 480 sec. After a dissociation time of 900 seconds (1500 seconds for scFv76-77), both flow cells surfaces were regenerated by flowing 10 mM Glycine-HCl, pH 1.5, at 30 µL/min for 30 sec and 10 mM NaOH at 30 µL/min for 30 sec. Double referenced sensorgrams were obtained by subtraction of blank-immobilized flow cell curves, as well as of zero concentration curves, from derivatized surface flow cell curves. Kinetic constants were obtained by BIAevaluation 3.2 software (Cytiva) fitting with a 1:1 binding model. For binning experiment by SPR, SARS-CoV-2 RBD mFc-tagged (Sino Biological, 10 µg/mL in acetate buffer, pH 5) was immobilized at 300 RU level on the surface of a flow cell of a Series S sensor chip CM5 (Cytiva) by classical amine coupling procedure, while another flow cell surface was blank-immobilized with ethanolamine to be used as control surface. Eight scFvs (76, 76-77, 76-58, 76-57, 76-55, 76-46, 86 and 5) were sequentially flowed on both surface at 100 nM in HBS-EP+ (Cytiva) with a contact time of 600 sec and a dissociation time of 60 sec for each protein, except for the last one (scFv5) that was let to dissociate for 240 sec. Then both flow cells surfaces were regenerated by flowing 10 mM glycine-HCl, pH 1.5, at 30 µL/min for 30 sec and 10 mM NaOH at 30 µL/min for 30 sec. Double referenced sensorgram was obtained by subtraction of blank-immobilized flow cell curve, as well as of sequentially zero concentration curve, from derivatized surface flow cell curve.

**HCS fluorescence imaging (Operetta)**

Human embryonic kidney HEK293T cells stably expressing the SARS-CoV-2 spike glycoprotein S1 (generated by transfection with plasmid DNA encoding for S1 protein and subsequent clonal selection), were seeded in 96-well microtiter plates (5 x $10^5$ cells/well) pretreated with polyethyleneimine (Sigma-Aldrich). After 24 h cells were washed with PBS, fixed with 4% formaldehyde, permeabilized with PBS/Tween (PBS, 0.2% Tween-20) and blocked with 2% BSA in PBS/Tween. Cells were then incubated with 50 µL/well of scFv at concentration of 10 µg/mL, for 1 h at 4 °C. After three washings with PBS/Tween, plates were treated with 10 µg/mL of a mouse anti-Flag M2-FITC (Sigma-Aldrich). Cells were ultimately counterstained with Draq5 (Axxora) and analyzed for fluorescence by means of the High Content Screening (HCS) system Operetta (Perkin Elmer).

**Flow Cytometry Analysis**

Human embryonic kidney HEK293T cells stably expressing the SARS-CoV-2 spike glycoprotein S1 (generated by transfection with plasmid DNA encoding for S1 protein and subsequent clonal selection), were washed with PBS and incubated with 360 nM scFv, for 1 h at 4 °C. After three washings with PBS, cells were treated with 10 µg/mL of a mouse anti-Flag M2-FITC (Sigma-Aldrich), washed again and analyzed for fluorescence by ATTUNE NxT acoustic focusing cytometer (Invitrogen-Thermo Fisher Scientific). Dead cells were rule out from analysis using propidium iodide staining (Sigma Aldrich, P4864).

**Microneutralization assay**

Neutralizing antibody titers were tested using a live-virus assay as follows. Decomplemented serum or scFv samples were pre-diluted in inoculation medium (DMEM, 2% fetal calf serum, 1% glutamine), followed by 9 serial dilutions in inoculation medium. Serial dilutions were then mixed 1:1 with 2000 TCID50/mL SARS-CoV-2 virus [strain 2019-nLOV/Italy INM1 or strain hCoV-19/USA/PHC658/2021 (B.1.617.2) WCCM] and incubated for 1 h at + 37 °C, 5% $CO_2$. Then 35 µL of each dilution/virus mix were applied in octuplicate to Vero E6 cells seeded at a density of $10^5$ cells/mL in a 96-well plate at day -1. After 1 h of incubation at 37 °C, 5% $CO_2$, 65 µL of culture medium (DMEM, 2% fetal calf serum, 1% glutamine) were added to each well. Plates were incubated for four days at 37 °C, 5% $CO_2$. After this period, cells were inspected for cytopathic effect (CPE) and the number of CPE-positive wells were recorded. The titer at which no CPE was observed in at least half of the octuplicates (50% micro-neutralization titer, MN50) was calculated according to Spearman-Kärber formula (*28*) for each sample.

**Viral Neutralization in Calu-3 cells**

To measure the SARS-CoV-2-neutralizing capability of antibodies and patient’s sera, a live SARS-CoV-2 assay was performed by measuring the viral load in human lung adenocarcinoma Calu-3 cells, 72 h after virus infection, by quantitative real-time reverse transcription-PCR (qRT-PCR). The experiments were carried out at the François Hyafil Research Institute (Oncodesign; Villebon-sur-Yvette, France). Calu-3 cells were seeded in 96-well plates in complete cell culture medium

(MEM + 1% pyruvate + 1% glutamine + 10% fetal bovine serum) and then infected, at a multiplicity of infection of 0.01, with three different SARS-CoV-2 virus strains: the SARS-CoV-2, 2019-nCoV/Japan/TY/WK-521/2020 strain, the Slovakia/SK-BMC5/2020 strain (endowed with the D614G mutation) and the Slovakia /SK-BMC-P1A/2021- B.1.1.7 variant (VOC 202012/01) strain (UK origin). Antibodies and a pool of patient's sera were assessed in two different approaches. In the first one, 1 h after infection the virus solution was discarded and replaced by a volume of growth medium containing each test compound, in a concentration ranging from 214 to 2.6 nM (or at dilutions from 1/20 to 1/320 for sera), in triplicate. In the second approach, test antibodies (in a concentration ranging from 214 to 0.097 nM) and sera (at the same dilutions as above) were premixed with the virus 1 h at room temperature and then added to cells. The plates were then transferred in a 37 °C incubator for 72 h. Finally, the cell culture supernatants were collected for viral RNA extraction with a Macherey Nagel Viral RNA kit and for viral RNA copy number detection (targeting a region in the IP2/IP4 gene) in a CFX384 Real-Time PCR Detection System (Bio-Rad Laboratories). Data were processed using GraphPad Prism software (V8.0) and the $IC_{50}$ values calculated using a four-parameter logistic curve fitting approach.

**Cell-cell fusion assay**

Human alveolar type II-like epithelial A549 cells, embryonic kidney 293T cells and monkey kidney Vero E6 cells were obtained from ATCC (Manassas, VA, USA). Cells were grown at 37 °C, 5% $CO_2$, in RPMI-1640 (A549 cells) or DMEM (Vero E6 and 293T cells) medium (Euroclone) supplemented with 10% fetal calf serum (FCS), 2 mM glutamine and antibiotics. Generation of A549 cells stably expressing the human ACE2 (hACE2) receptor (A549-hACE2 cells) has been described previously (*29*). The C-terminal Flag-tagged pCMV3-2019-nCoV-spike (S1+S2)-long-Flag (SARS-2, Wuhan) and the pCMV3-SARS-CoV-2-spike (S1+S2)-long (B.1.617.2) (Delta) vectors were obtained from Sino Biological. Flag-tagged pCAGGS-SARS2-S-G614 (Addgene plasmid #156421) (D614G) vector was a gift from Hyeryun Choe and Michael Farzan (*30*). The pCMV-GFP vector was obtained from Clontech. Transfections were performed using Lipofectamine 2000 (Invitrogen, Thermo Scientific), according to the manufacturer's instructions. In the donor-target cell fusion assay, 293T cells were co-transfected with plasmids encoding the Flag-tagged SARS-CoV-2 Wuhan S-protein, the Flag-tagged D614G S protein or the spike protein from the SARS-CoV-2 Delta variant, and GFP. 293T cells, co-transfected with the GFP-encoding plasmid and the empty vector, were used as negative control. At 40 h post transfection cells were detached with trypsin (0.25%) and incubated with different concentrations of scFv76 or scFv5 antibodies at 37 °C. After 30 or 60 min, cells were overlaid on an 80% confluent monolayer of target cells [Vero E6 (naturally expressing ACE2 receptors on the membrane surface) or A549-hACE2 (stably transfected with the hACE2 encoding plasmid) cells] at a ratio of approximately one S expressing cell to three receptor expressing cells (*31*). After 4 h nuclei were stained with Hoechst. Transmission and fluorescence images were taken using a ZEISS Axio Observer inverted microscope. To measure the extent of cell-cell fusion, the GFP area was delimited, measured using ImageJ software (*32*), divided by the total cell area and expressed as percentage relative to control. Images shown in all figures are representative of at least five random fields (scale-bars are indicated). Statistical analysis was performed using one-way ANOVA test (Prism 6.0 software, GraphPad). All experiments were done in duplicate and repeated at least twice.

**Generation of SARS-CoV-2 S-pseudoviruses**

Embryonic kidney 293T cells (ATCC) were grown at 37 °C, 5% $CO_2$, in DMEM medium (Euroclone) supplemented with 10% FCS, 2 mM glutamine and antibiotics. The C-terminal Flag-tagged pCMV3-2019-nCoV-spike (S1+S2)-long-Flag (Wuhan) and pCAGGS-SARS2-S-G614 (Addgene plasmid #156421) (D614G), and the pCMV3-SARS-CoV-2-spike (S1+S2)-long (B.1.617.2) (Delta) vectors are described above. The mutants N501Y and K417N+E484K+N501Y were constructed on the

Wuhan plasmid; mutations were generated and sequence-validated by Bio-Fab Research (Rome, IT). The C-terminal Flag-tagged pReceiver-M39 UK B.1.1.7 (EX-CoV242-M39) (Alpha), South Africa B.1.351 (EX-CoV244-M39-GS) (Beta) and Brazil B.1.1.28 (EX-CoV245-M39-GS) (Gamma) vectors were obtained from GeneCopoeia.
Transfections were performed using Lipofectamine 2000 (Invitrogen, Thermo Scientific). For production of SARS-CoV-2 S-pseudotyped lentiviral particles (LV) carrying a firefly luciferase reporter gene, 293T cells were co-transfected with the Wuhan plasmid, the D614G plasmid or plasmids encoding spike protein variants (Alpha, Beta, Gamma, Delta), the pCMVR 8.74 lentiviral packaging plasmid (a kind gift from R. Piva, University of Turin, Italy) and the pLenti CMV Puro LUC plasmid (Addgene plasmid #17477), a gift from Eric Campeau and Paul Kaufman (*33*). At 48 h after transfection, supernatants containing the viral particles were collected, filtered through 0.45 µm membranes and frozen at -80 °C until use. Spike-pseudotyped LV titers were determined assessing the number of viral RNA genomes/mL using RT-qPCR. Briefly, viral RNAs extracted with TRIzol-LS Reagent (Invitrogen, Thermo Scientific) were digested with TURBO DNase (Invitrogen, Thermo Scientific) at 37 °C for 15 min. Reverse transcription of viral RNA (5 µL) was performed using PrimeScript RT Reagent Kit (Takara). RT-qPCR analysis was carried out with a CFX-96 Touch Real-Time PCR Detection System (Bio-Rad) using the SensiFAST SYBR® kit (Bioline) and primers specific for the luciferase-reporter gene (Luc F: 5’-CATTCCGGATACTGCGATTT-3’; Luc R: 5’-GGCGAAGAAGGAGAATAGGG-3’). To generate a standard curve, a 451 bp (position 544-994) luciferase-RNA fragment was transcribed *in vitro* using MEGAshortscript T7 Transcription Kit (Invitrogen, Thermo Scientific). Pseudovirus particles were adjusted to the same titer (RNA copies/mL) in all neutralization experiments.
For *in vivo* experiments, LV pseudotyped with the SARS-CoV-2 D614G-S variant were produced as described above, concentrated using PEG Virus Precipitation Kit (Abcam), resuspended in Abcam VR (virus re-suspension) solution and then frozen at -80 °C until use.

**SARS-CoV-2 S-pseudovirus neutralization assay**

Human Caco-2 hACE2 cells, stably expressing the hACE2 receptor, were grown at 37°C, 5% $CO_2$, in DMEM medium (Euroclone) supplemented with 10% FCS, 2 mM glutamine and antibiotics. Generation of Caco-2 hACE2 cells has been described previously (*29*). Caco-2 hACE2 cells were plated at a density of $3x10^4$ cells/well in white clear-bottom 96-well plates (Costar). After 24 h, 100 µL of DMEM containing 5% FCS and 8 µg/mL polybrene were added to each well, and cells were incubated for 30 min. Serial (1:3) dilutions (range: from 1 µg/mL to 0.004 µg/mL final concentration) of scFvs were prepared in DMEM containing 5% FCS. Each dilution was tested in duplicate on each plate. Aliquots (60 µL) of the LV suspension were added to an equal amount of each antibody dilution (60 µL) and incubated for 1 h at 37 °C. Pseudovirus:scFv mixtures (50 µL) were then added to Caco-2 hACE2 cells after medium removal. Additional 50 µL of DMEM containing 5% FCS were added to each well after 24 h. Luciferase activity (Relative Luciferase Units or RLU) was detected at 72 h post-infection by using Bright-Glo Luciferase Assay System Kit (Promega) in a Microplate Luminometer (Wallac-Perkin Elmer).

**Epitope mapping**

Shotgun Mutagenesis epitope mapping services were provided by Integral Molecular (Philadelphia, PA) as described in Davidson and Doranz, 2014 (*34*). Briefly, a mutation library of the target protein was created by high-throughput, site-directed mutagenesis. Each residue was individually mutated to alanine, with alanine codons mutated to serine. The mutant library was arrayed in 384-well microplates and transiently transfected into HEK 293-T cells. Following transfection, cells were incubated with the indicated antibodies at concentrations pre-determined using an independent immunofluorescence titration curve on wild type protein. Test antibodies were detected using an Alexa Fluor 488-conjugated secondary antibody and mean cellular fluorescence was determined using Intellicyt iQue flow cytometry platform. Mutated residues were identified as being critical to the test antibody epitope if they did not support the reactivity of the test antibody but did support the reactivity of the reference MAb. This counterscreen strategy facilitates the exclusion of mutants that are locally misfolded or that have an expression defect.

### Biochemical analyses

SDS-PAGE was performed by standard methods (*35*). SEC-HPLC analyses were performed on an ALLIANCE HPLC system equipped with a 2998 Photodiode Array detector (Waters). Antibodies were loaded on a TSKgel G3000 SWXL 30 cm x 7.8 mm column (Tosoh Bioscience) at a concentration of 3 µg in 100 µL PBS and eluted with running buffer (50 mM phosphate buffer, 150 mM NaCl, pH 7.2), or running buffer/acetonitrile mixture (90:10 v/v), at a flow rate of 0.75 mL/min. Peaks were detected at 280 nm. For some analyses, antibodies were loaded on a BioSep™ 5µM SEC-S2000 145Å 300 x 7,8mm column (PHENOMENEX), and eluted with running buffer or running buffer/acetonitrile mixture (90:10 v/v), at a flow rate of 1mL/min. Detection by 280 nM absorbance.

SEC-MS analysis was performed on an UHPLC Ultimate 3000SD (Thermo Fisher) coupled to a Q-Exactive Plus mass spectrometer (Thermo Fisher) operating in electrospray ionization in positive ion mode. The analysis was performed on a column Mab Pac SEC-1 (2.1 mm x 100 mm, Thermo Fisher), maintained at 25 °C using 20 mM ammonium formate. The mass spectra were recorded from m/z 1500 to m/z 8000 in native conditions.

The primary amino acid sequences were confirmed by peptide mass fingerprint analysis. Samples were reduced, alkylated and digested with trypsin. The resulting peptide mixtures were analyzed by RP-UHPLC-MS/MS. The samples were injected on a column Acquity UPLC Waters CSH C18 130Å (1 mm x 100 mm, 1.7 µm, Waters). The column oven was maintained at 55 °C, the analysis was carried using a gradient elution (phase A: 0.1% formic acid in water; phase B: 0.1% formic acid in acetonitrile). The flow rate was maintained at 0.1 mL/min. The mass spectra were acquired in "data dependent scan" mode, able to acquire both the full mass spectra in high resolution and to isolate and fragment the ten ions with highest intensity present in the mass spectrum. The raw data obtained were analyzed using the Biopharma Finder 2.1 software (Thermo Fisher). The elaboration process consisted in the comparison between the peak list obtained "in silico" considering an expected amino acid sequence of a certain protein and fixed or variable modifications (carboamidomethylation, oxidation etc.) and the experimental data.

### Circular Dichroism (CD)

CD experiments were performed on a Jasco J-815 spectropolarimeter equipped with a PTC-423S/15 Peltier temperature controller. Cells with 0.1 cm path length and scFv protein concentrations of 0.2–0.9 mg/mL were used to record CD spectra between 200 and 250 nm, with a 2 nm bandwidth, and a scan rate of 5 or 20 nm/min. Spectra were recorded at 20 and 90 °C. Spectra were signal-averaged over five scans, baseline corrected by subtracting a buffer spectrum, and smoothed using the means-movement function before conversion to molar ellipticity. Spectra were analyzed for secondary structure composition using the BeStSel method (*20*).

### *In vivo* models of infection with SARS-CoV-2 or SARS-CoV-2 S-pseudotyped virus

Animal studies were performed in accordance with the European Directive 2010/63/EU on the protection of animals used for scientific purposes, applied in Italy by the Legislative Decree 4 March

2014, n. 26. This study was included in a main research project approved by Italian Department of Health. Female transgenic mice (K18-hACE2), aged 4-6 weeks, were infected via the intranasal route (10 μL/nostril) with a LV pseudotyped with the SARS-CoV-2 D614G-S variant expressing luciferase (Lenti-LUC-spike D614G), generated as described above. A luciferase-expressing lentivirus was used (10 μL/nostril) as a negative control. Two h before and 4 h after infection, scFv76 (3.7 mg/mL in PBS) and scFv76-58 (2.6 mg/mL in PBS) antibodies were intranasally administered (10 μL/nostril and 14 μL/nostril, respectively) to mice (n=12/group). An unrelated scFv (2.6 mg/mL in PBS) was also administered (14 μL/nostril) to additional mice (n=12) in the same experimental conditions as above. At three different time points (48 h, 72 h and 96 h post infection) Lenti-LUC-spike D614G bioluminescence was quantitatively measured through the IVIS200 optical imaging system (Perkin Elmer) for each mouse. Total body bioluminescence imaging was performed on mice before euthanasia, then explanted lungs and nasal turbinates were imaged. In vivo scFv testing against an authentic SARS-CoV-2 strain was performed by using a Syrian hamster model at the François Hyafil Research Institute (Oncodesign; Villebon-sur-Yvette, France). Animal housing and experimental procedures were conducted according to the French and European Regulations and the National Research Council Guide for the Care and Use of Laboratory Animals. SARS-CoV-2 strain "Slovakia/SK-BMC5/2020", originally provided by the European Virus Archive global (EVAg) with D614G mutation, produced and tittered by Oncodesign on Vero E6/TMPRSS2 cells, was used for hamster infection. Female golden Syrian 7-week old hamsters, purchased from Janvier Labs, were anesthetized with isoflurane and inoculated intranasally with 70 µL (35 µL/nostril) containing a dose of $10^5$ pfu SARS-CoV-2. ScFv76 (1 mg/mL in D-PBS) was administered by intranasal route to isoflurane-anesthetized animals (n=5), under a volume of 70 µL (35 µL/nostril), 2 h before infection and then once daily for two consecutive days post infection. Hamsters were monitored for appearance, behavior and weight and ultimately were euthanized at day 4 post infection by intraperitoneal injection of a cocktail of Zoletil (30 mg/kg) and Xylazine (10 mg/kg) followed by gentle cervical dislocation. Thoracotomy was then performed before tissue collection and lung necropsy.

#### *In vivo* nebulization of 76clAbs

Animal studies were performed in accordance with the European Directive 2010/63/EU on the protection of animals used for scientific purposes, applied in Italy by the Legislative Decree 4 March 2014, n. 26. This study was included in a research project approved by Italian Department of Health. Female BALB/c OlaHsd mice, purchased from Envigo, were nebulized with scFv antibodies at various concentrations (ranging from 1 to 5 mg/mL) in DPBS, and at different duration of nebulization exposure, by using a home-made equipment, based on the Aerogen Pro (Aerogen) mesh nebulizer and a nose-only inhalation chamber, suitable for delivering the nebulized antibodies contemporarily up to 8 mice. At different time points post-nebulization mice were euthanized and lungs explanted and then fixed by 4% paraformaldehyde for IHC analyses or snap-frozen (in liquid nitrogen) and stored at -80 °C until further analysis.

For assessing the recovery of functional scFv antibody, the snap-frozen lungs were weighed, transferred to different tubes on ice in the presence of 1x Lysis buffer (Cell Signaling), already containing different protease inhibitors and supplemented with 1 mM PMSF. The lung tissues were homogenized at 4°C for 20 sec through a rotor-stator homogenizer, then incubated on ice for 15 min and sonicated for 30 sec by a suitable ultrasonic bath. Lung homogenates were centrifuged at 14,000 × g for 10 minutes at 4°C, and then supernatants were transferred to clean microcentrifuge tubes and frozen on dry ice and stored at -80°C. Total protein concentrations in the lung tissue homogenates were determined using a classical Bradford assay. For testing whether the nebulized antibodies conserved their functional properties, aliquots of lung tissue homogenates were diluted in PBS buffer containing tween and BSA and then assessed by ELISA for the binding to SARS-CoV-2 RBD as previously described.

### Immunohistochemistry

Immunohistochemistry staining to detect the scFv antibodies in mice lungs after nebulization was performed with the His-Tag antibody Rabbit monoclonal antibody, clone RM146 (cat. SAB5600227, from Sigma-Aldrich). Slides were deparaffinized and treated with 3% $H_2O_2$ in $H_2O$ to quench endogenous peroxidase activity. ScFv antibodies staining was performed at 4 °C ON, followed by exposure to biotinylated-anti-rabbit secondary antibody for 60 min at room temperature. Diaminobenzidine (DAB) was used as the chromogen. Microscopic observation was performed using a Nikon Eclipse 80i microscope equipped with a DXM1200F Microscope Camera. Cross-reactivity of scFv76 with human normal tissues was assessed by using Tissue microarrays (TMAs), obtained from US Biomax Inc. (cat. MN0341). To this aim, slides were treated as described above and then incubated with scFv (10 µg/mL) overnight at 4°C, followed by the same procedures described before. An anti-hPDL1 scFv was used as positive control.

**Table S1. Viral neutralization by COVID-19 convalescent sera**

| | MN50 | | |
|---|---|---|---|
| **CS6** | 1/100 | | |
| **CS9** | 1/100 | | |
| **CS10** | 1/100 | | |
| **CS8** | | 1/50 | |
| **CS1** | | 1/50 | |
| **CS4** | | 1/50 | |
| **CS2** | | | Undetected |
| **CS5** | | | Undetected |
| **CS3** | | | Undetected |
| **NS pool** | | | Undetected |

Neutralizing activity against SARS-CoV-2 (Wuhan strain) of sera from 10 COVID-19 convalescent patients by micro-neutralization assay in Vero E6 cells. Sera were decomplemented for 30 min at 56 °C and then added to cells after 2-fold dilutions ranging from 1/50 up to 1/800 (8 replicates). Cytopathogenic effect (CPE) was measured after 4 days cultivation. Data are expressed as the titer (dilution) at which no CPE was observed in at least half of wells (50% micro-neutralization titer, MN50) calculated according to the Spearman-Kärber formula.

**Table S2. Sequence alignment CDR3 of heavy and light chain and germline usage of anti-SARS-CoV-2 human scFv antibodies**

| ScFv | VH CDR3 | Germline | VL CDR3 | Germline |
|---|---|---|---|---|
| **76** | ARDLSVAGAFDI | IGVH3-53 | QQYGSSP_YT | IGVK3-20 |
| **76-46** | ARDLSVAGAFDI | IGVH3-53 | QQYGSSP_IT | IGVK3-20 |
| **76-55** | ARDLSVAGAFDI | IGVH3-53 | QQYVRSP_PT | IGVK3-15 |
| **76-57** | ARDLSVAGAFDI | IGVH3-53 | QHRD_____I | IGVK1-9 |
| **76-58** | ARDLSVAGAFDI | IGVH3-53 | QQYGSSPRIT | IGVK3-20 |
| **76-77** | ARDLSVAGAFDI | IGVH3-53 | QEYGSSPRVT | IGVK3-20 |
| **5** | ATGGAVPHVTGAFDI | IGVH1-46 | NSYTRFSTGV | IGVL2-14 |
| **86** | ATGGAVPHVTGAFDI | IGVH1-46 | QQRSNWPPGFT | IGVK3-11 |

Sequence alignment was performed by IGBLAST tool https://www.ncbi.nlm.nih.gov/igblast/

**Table S3. Binding of anti-SARS-CoV-2 scFv antibodies to spike and RBD mutants**

| scFv | S1WT | S1D614G | S1alpha | S1beta | S1delta |
|---|---|---|---|---|---|
| **76** | 0.25(0.11) | 0.14(0.04) | 0.21(0.14) | 0.54(0.29) | 0.18(0.05) |
| **76-46** | 0.29(0.11) | 0.14(0.04) | 0.29(0.21) | 0.61(0.36) | 0.30(0.03) |
| **76-55** | 0.57(0.40) | 0.25(0.14) | 0.43(0.04) | >10xWT | 0.52(0.08) |
| **76-57** | 0.51(0.36) | 0.25(0.15) | 0.44(0.04) | 1.34(0.76) | 0.56(0.03) |
| **76-58** | 0.39(0.21) | 0.25(0.11) | 0.32(0.21) | 0.89(0.46) | 0.27(0.03) |
| **76-77** | 0.25(0.04) | 0.18(0.04) | 0.21(0.14) | 0.43(0.14) | 0.21(0.00) |
| **5** | 2.91(1.49) | 1.06(0.32) | 1.70(1.74) | 4.79(4.58) | 1.79(0.13) |
| **86** | 1.77(1.17) | 0.71(0.28) | 1.03(0.78) | 3.16(2.45) | 1.14(0.10) |

| scFv | WT | N501Y | V483A | I472V | Y453F | A475V | E484K | N439K | A520S | L452R | K417N | E484Q | L452R/E484Q | L452R/T478K | T478K |
|---|---|---|---|---|---|---|---|---|---|---|---|---|---|---|---|
| **76** | 0.11(0.04) | 0.11(0.04) | 0.21(0.11) | 0.21(0.11) | 0.11(0.04) | 0.43(0.14) | 0.54(0.04) | 0.25(0.21) | 0.14(0.04) | 0.11(0.04) | 0.14(0.04) | 0.23(0.03) | 0.21(0.00) | 0.20(0.03) | 0.16(0.03) |
| **76-46** | 0.18(0.07) | 0.18(0.00) | 0.32(0.18) | 0.36(0.18) | 0.18(0.07) | 0.54(0.29) | 0.64(0.04) | 0.36(0.25) | 0.32(0.11) | 0.29(0.14) | 0.18(0.04) | 0.30(0.03) | 0.30(0.03) | 0.34(0.03) | 0.30(0.03) |
| **76-55** | 0.14(0.04) | 0.32(0.22) | 0.79(1.01) | 0.83(1.08) | 0.32(0.25) | >10xWT | >10xWT | >10xWT | 0.32(0.22) | 0.29((0.18) | 0.32(0.04) | 0.45(0.03) | 0.65(0.05) | 0.59(0.03) | 0.49(0.03) |
| **76-57** | 0.18(0.07) | 0.33(0.22) | 0.58(0.65) | 0.62(0.73) | 0.33(0.22) | 1.12(1.12) | 1.12(0.58) | 1.34(1.45) | 0.33(0.22) | 0.29(0.22) | 0.36(0.04) | 0.71(0.03) | 0.74(0.03) | 0.56(0.03) | 0.54(0.00) |
| **76-58** | 0.21(0.07) | 0.21(0.04) | 0.39(0.28) | 0.60(0.64) | 0.21(0.04) | 0.60(0.21) | 1.64(0.64) | 0.85(0.82) | 0.25(0.11) | 0.25(0.07) | 0.18(0.00) | 0.48(0.03) | 0.48(0.03) | 0.27(0.03) | 0.23(0.03) |
| **76-77** | 0.11(0.04) | 0.11(0.00) | 0.28(0.11) | 0.25(0.14) | 0.14(0.04) | 0.32(0.21) | 0.43(0.14) | 0.28(0.21) | 0.21(0.07) | 0.18(0.04) | 0.14(0.00) | 0.27(0.03) | 0.27(0.03) | 0.23(0.03) | 0.21(0.00) |
| **5** | 1.06(0.32) | 3.19(1.92) | >10xWT | >10xWT | 3.09(0.75) | >10xWT | >10xWT | >10xWT | 1.06(0.50) | 1.77(0.92) | 1.99(0.00) | 8.08(0.38) | 4.58(0.05) | 3.23(0.50) | 3.14(0.58) |
| **86** | 0.60(0.25) | 1.38(0.18) | >10xWT | >10xWT | 1.42(0.21) | >10xWT | >10xWT | >10xWT | 0.75(0.50) | 0.85(0.53) | 0.96(0.11) | 2.59(0.05) | 1.37(0.03) | 1.51(0.23) | 1.30(0.08) |

Binding of scFv antibodies to SARS-CoV-2 spike (**top**) and RBD (**bottom**) variants measured by ELISA. Data are expressed as nM concentration to have 1.0 OD (λ 450 nm) and are the average (±SD) of 2-3 independent experiments. S1(WT): Wuhan strain; S1(D614G) present in all variants; S1alpha: mutations HV69-70d, Y144d, N501Y, A570D, D614G, P681H as in Alpha (B.1.1.7) VBM; S1beta: mutations K417N, E484K, N501Y, D614G as in Beta (B.1.351) VBM and Gamma (P.1) VBM (K417T); S1delta: mutations T19R, G142D, E156G, 157-158del, L452R, T478K, D614G, P681R as in Delta (B.1.617.2) VOC. Double RBD mutations L452R/E484Q as in Kappa (B.1.617.1) VBM, or L452R/T478K as in Delta (B.1.617.2) VOC. Boxed numbers indicate fold increase >10x versus WT.

**Table S4. Affinity of anti-SARS-CoV-2 scFv antibodies for mutated RBD proteins by Surface Plasmon Resonance**

**scFv 76**

| RBD | $K_a$ ($10^5$ $M^{-1}$ $s^{-1}$) | $k_d$ ($10^{-5}$ $s^{-1}$) | $K_D$ (nM) |
|---|---|---|---|
| Wt | 1.2 | 7.3 | 0.6 |
| N439K | 0.9 | 8.5 | 1.0 |
| L452R | 0.9 | 8.0 | 0.9 |
| Y453F | 1.0 | 11.8 | 1.2 |
| I472V | 1.0 | 8.5 | 0.9 |
| A475V | 1.0 | 57.0 | 5.5 |
| V483A | 1.4 | 9.6 | 0.7 |
| E484K | 0.9 | 10.7 | 1.2 |
| N501Y | 0.8 | 4.7 | 0.6 |
| A520S | 1.4 | 7.9 | 0.6 |
| K417N | 1.6 | 42.8 | 2.7 |
| L452R, E484Q | 1.3 | 12.7 | 1.0 |
| L452R, T478K | 0.8 | 7.3 | 0.9 |

**scFv 76-77**

| RBD | $K_a$ ($10^5$ $M^{-1}$ $s^{-1}$) | $k_d$ ($10^{-5}$ $s^{-1}$) | $K_D$ (nM) |
|---|---|---|---|
| wt | 1.2 | 4.2 | 0.4 |
| N439K | 0.9 | 6.6 | 0.7 |
| L452R | 1.0 | 5.6 | 0.6 |
| Y453F | 1.0 | 7.2 | 0.7 |
| I472V | 1.0 | 7.7 | 0.8 |
| A475V | 1.0 | 37.0 | 3.7 |
| V483A | 1.0 | 5.5 | 0.6 |
| E484K | 0.7 | 6.3 | 1.0 |
| N501Y | 1.0 | 3.1 | 0.3 |
| A520S | 1.0 | 4.8 | 0.5 |
| K417N | 1.2 | 26.7 | 2.3 |
| L452R, E484Q | 0.8 | 8.7 | 1.0 |
| L452R, T478K | 0.8 | 5.6 | 0.7 |

Continues….

Continues Table S4

**scFv 76-55**

| RBD | $K_a$ ($10^5$ $M^{-1}$ $s^{-1}$) | $k_d$ ($10^{-5}$ $s^{-1}$) | $K_D$ (nM) |
|---|---|---|---|
| wt | 1.3 | 45.2 | 3.6 |
| N439K | 1.0 | 36.7 | 3.5 |
| L452R | 1.1 | 47.8 | 4.3 |
| Y453F | 1.0 | 55.1 | 5.4 |
| I472V | 1.3 | 48.7 | 3.6 |
| A475V | 1.0 | 84.6 | 8.5 |
| V483A | 1.2 | 63.6 | 5.4 |
| E484K | 0.9 | 74.5 | 8.7 |
| N501Y | 1.0 | 77.0 | 7.4 |
| A520S | 1.2 | 56.0 | 4.6 |
| K417N | 1.7 | 72.8 | 4.3 |
| L452R, E484Q | 1.6 | 51.1 | 3.2 |
| L452R, T478K | 0.9 | 58.6 | 6.7 |

**scFv 76-57**

| RBD | $K_a$ ($10^5$ $M^{-1}$ $s^{-1}$) | $k_d$ ($10^{-5}$ $s^{-1}$) | $K_D$ (nM) |
|---|---|---|---|
| wt | 1.2 | 7.3 | 0.6 |
| N439K | 0.9 | 11.5 | 1.3 |
| L452R | 1.0 | 7.6 | 0.8 |
| Y453F | 1.1 | 5.3 | 0.5 |
| I472V | 1.0 | 11.3 | 1.2 |
| A475V | 1.3 | 52.7 | 4.1 |
| V483A | 1.1 | 12.3 | 1.2 |
| E484K | 0.7 | 12.2 | 1.7 |
| N501Y | 1.1 | 14.4 | 1.3 |
| A520S | 1.1 | 9.4 | 0.9 |
| K417N | 1.5 | 41.0 | 2.8 |
| L452R, E484Q | 1.1 | 14.0 | 1.2 |
| L452R, T478K | 0.7 | 8.3 | 1.2 |

Continues….

Continues Table S4

**scFv 76-58**

| RBD | $K_a$ ($10^5$ $M^{-1}$ $s^{-1}$) | $k_d$ ($10^{-5}$ $s^{-1}$) | $K_D$ (nM) |
|---|---|---|---|
| wt | 2.8 | 41.1 | 1.5 |
| N439K | 2.4 | 39.6 | 1.6 |
| L452R | 2.5 | 41.6 | 1.7 |
| Y453F | 2.2 | 66.7 | 3.0 |
| I472V | 2.8 | 41.8 | 1.5 |
| A475V | 2.3 | 52.2 | 2.2 |
| V483A | 2.6 | 52.3 | 2.0 |
| E484K | 1.3 | 44.8 | 3.5 |
| N501Y | 2.7 | 38.4 | 1.4 |
| A520S | 2.6 | 46.2 | 1.7 |
| K417N | 2.4 | 34.6 | 1.5 |
| L452R, E484Q | 1.4 | 35.5 | 2.6 |
| L452R, T478K | 1.5 | 42.9 | 2.8 |

**scFv 76-46**

| RBD | $K_a$ ($10^5$ $M^{-1}$ $s^{-1}$) | $k_d$ ($10^{-5}$ $s^{-1}$) | $K_D$ (nM) |
|---|---|---|---|
| wt | 1.4 | 4.0 | 0.3 |
| N439K | 1.0 | 3.2 | 0.3 |
| L452R | 1.3 | 4.2 | 0.3 |
| Y453F | 1.2 | 4.4 | 0.4 |
| I472V | 1.2 | 7.3 | 0.6 |
| A475V | 1.6 | 70.1 | 4.5 |
| V483A | 1.3 | 4.9 | 0.4 |
| E484K | 0.9 | 4.9 | 0.6 |
| N501Y | 1.1 | 9.1 | 0.9 |
| A520S | 1.4 | 4.2 | 0.3 |
| K417N | 1.8 | 75.1 | 4.2 |
| L452R, E484Q | 1.2 | 9.1 | 0.7 |
| L452R, T478K | 0.8 | 2.5 | 0.3 |

Continues….

Continues Table S4

**scFv 5**

| **RBD** | ***$K_a$*** **($10^5$ $M^{-1}$ $s^{-1}$)** | ***$k_d$*** **($10^{-5}$ $s^{-1}$)** | ***$K_D$*** **(nM)** |
|---|---|---|---|
| wt | 0.3 | 19.1 | 6.2 |
| N439K | 0.3 | 16.3 | 5.2 |
| L452R | 0.3 | 14.6 | 4.9 |
| Y453F | 0.3 | 16.8 | 5.4 |
| I472V | 0.3 | 19.3 | 6.8 |
| A475V | 0.3 | 15.1 | 5.0 |
| V483A | 0.3 | 18.1 | 6.1 |
| E484K | 0.3 | 17.6 | 6.0 |
| N501Y | 0.3 | 19.1 | 6.7 |
| A520S | 0.4 | 12.3 | 3.0 |
| K417N | 0.3 | 14.6 | 5.4 |
| L452R, E484Q | 0.3 | 20.4 | 6.4 |
| L452R, T478K | 0.4 | 15.9 | 4.5 |

**scFv 86**

| **RBD** | ***$K_a$*** **($10^5$ $M^{-1}$ $s^{-1}$)** | ***$k_d$*** **($10^{-5}$ $s^{-1}$)** | ***$K_D$*** **(nM)** |
|---|---|---|---|
| wt | 0.5 | 13.1 | 2.8 |
| N439K | 0.5 | 9.9 | 2.0 |
| L452R | 0.5 | 9.6 | 1.9 |
| Y453F | 0.5 | 9.6 | 1.9 |
| I472V | 0.5 | 11.4 | 2.3 |
| A475V | 0.5 | 8.3 | 1.6 |
| V483A | 0.5 | 10.0 | 2.1 |
| E484K | 0.5 | 10.0 | 2.1 |
| N501Y | 0.4 | 10.3 | 2.4 |
| A520S | 0.6 | 8.7 | 1.5 |
| K417N | 0.5 | 9.3 | 1.8 |
| L452R, E484Q | 0.6 | 9.1 | 1.6 |
| L452R, T478K | 0.4 | 10.4 | 2.5 |

**Table S5. Inhibitory activity ($IC_{50}$) of anti-SARS-CoV-2 scFv antibodies on spike or RBD binding to human ACE2**

| scFv | S1WT | S1D614G | S1alpha | S1beta | S1delta |
|---|---|---|---|---|---|
| 76 | 0.62(0.11) | 1.11(0.07) | 1.07(0.21) | 1.84(0.14) | 1.64(0.25) |
| 76-46 | 0.54(0.05) | 1.23(0.12) | 1.72(0.25) | 1.94(0.09) | 1.86(0.25) |
| 76-55 | 1.87(0.15) | 2.65(0.15) | 2.94(0.37) | >40 | 3.02(0.18) |
| 76-57 | 1.61(0.24) | 2.49(0.16) | 2.56(0.13) | 2.63(0.34) | 3.08(0.25) |
| 76-58 | 0.77(0.12) | 1.44(0.21) | 1.73(0.09) | 2.74(0.21) | 2.49(0.04) |
| 76-77 | 0.36(0.08) | 0.72(0.15) | 0.90(0.13) | 1.62(0.16) | 1.95(0.04) |

| scFv | WT | N501Y | V483A | I472V | Y453F | A475V | E484K | N439K | A520S | L452R | K417N | E484Q | L452R/E484Q | L452R/T478K | T478K |
|---|---|---|---|---|---|---|---|---|---|---|---|---|---|---|---|
| 76 | 0.67(0.10) | 1.20(0.14) | 1.29(0.11) | 1.18(0.17) | 2.16(0.14) | 1.74(0.24) | 0.88(0.16) | 1.37(0.11) | 1.04(0.07) | 0.91(0.07) | 1.63(0.56) | 2.07(0.32) | 2.28(0.32) | 2.03(0.25) | 1.96(0.00) |
| 76-46 | 1.22(0.21) | 1.55(0.08) | 2.20(0.12) | 1.44(0.31) | 1.93(0.13) | 1.92(0.30) | 1.15(0.38) | 1.11(0.06) | 1.31(0.12) | 1.16(0.34) | 1.79(0.58) | 2.58(0.18) | 2.25(0.04) | 2.15(0.07) | 2.36(0.21) |
| 76-55 | 2.03(0.32) | 2.71(0.09) | 2.98(0.16) | 1.99(0.02) | 3.57(0.38) | 3.45(0.40) | 1.77(0.14) | 2.09(0.34) | 2.19(0.27) | 1.84(0.31) | 2.59(0.36) | 4.49(0.00) | 5.25(0.36) | 2.98(0.32) | 3.16(0.07) |
| 76-57 | 2.09(0.15) | 2.59(0.25) | 2.49(0.17) | 2.42(0.25) | 2.83(0.19) | 4.30(1.11) | 2.00(0.04) | 2.05(0.33) | 2.18(0.24) | 2.27(0.08) | 1.98(0.31) | 6.64(0.00) | 3.99(1.63) | 3.05(0.18) | 3.12(0.18) |
| 76-58 | 1.45(0.21) | 1.74(0.12) | 1.48(0.24) | 1.54(0.12) | 2.31(0.15) | 3.91(1.00) | 2.06(0.11) | 1.42(0.27) | 1.50(0.17) | 1.74(0.15) | 1.76(0.59) | 2.60(0.07) | 3.67(1.25) | 2.67(0.28) | 2.49(0.39) |
| 76-77 | 0.58(0.04) | 0.78(0.16) | 0.77(0.21) | 1.19(0.07) | 1.36(0.07) | 1.69(0.19) | 0.97(0.16) | 1.14(0.32) | 0.79(0.08) | 0.88(0.08) | 1.67(0.75) | 1.74(0.28) | 1.63(0.21) | 2.20(0.11) | 1.81(0.18) |

Competition of scFv antibodies in spike (**top**) or RBD (**bottom**) binding to human ACE2 measured by ELISA. $IC_{50}$ values (expressed as nM concentration) are the average (±SE) from 2-4 independent experiments. Boxed number indicates concentration >40 nM. S1(WT): Wuhan strain; S1(D614G) present in all variants; S1alpha: mutations HV69-70d, Y144d, N501Y, A570D, D614G, P681H as in Alpha (B.1.1.7) VBM; S1beta: mutations K417N, E484K, N501Y, D614G as in Beta (B.1.351) VBM and Gamma (P.1) VBM (K417T); S1delta: mutations T19R, G142D, E156G, 157-158del, L452R, T478K, D614G, P681R as in Delta (B.1.617.2) VOC. Double RBD mutations L452R/E484Q as in Kappa (B.1.617.1) VBM, or L452R/T478K as in Delta (B.1.617.2) VOC. ScFv5 and scFv86 antibodies did not interfere with spike or RBD binding to ACE2 at concentrations ≥40 nM.

**Table S6. Virus neutralization ($IC_{50}$) by anti-SARS-CoV-2 scFv antibodies in Calu-3 cells**

| | | $IC_{50}$ (nM) | |
|---|---|---|---|
| **scFv** | **Virus** | **Added to cells 1h after infection** | **Pre-incubated 1h with the virus before cell infection** |
| **76** | WT | 12.02 | <0.1 |
| | D614G | 13.02 | 1.13 |
| | Alpha VBM | 5.23 | 2.18 |
| **76-46** | WT | 19.35 | <0.1 |
| | D614G | 21.27 | 1.53 |
| | Alpha VBM | ND | ND |
| **76-58** | WT | 4.56 | 0.29 |
| | D614G | 4.03 | 0.67 |
| | Alpha VBM | 3.79 | 1.40 |
| **UR** | WT | >1,400 | >1,400 |
| | D614G | >1,400 | >1,400 |
| | Alpha VBM | >1,400 | >1,400 |
| **CS*** | WT | >1/200 | >1/10,000 |
| | D614G | >1/200 | >1/10,000 |
| | Alpha VBM | ND | ND |

Serially diluted (3-fold) antibodies were put in contact for 1 h with SARS-CoV-2 WT (Wuhan strain), SARS-CoV-2 variant with D614G mutation or SARS-CoV-2 Alpha variant, and then added to Calu-3 cell culture, or directly added to the cells 1 h after viral infection. Quantification of viral load was done by RT-qPCR 72 h after infection. $IC_{50}$ values (expressed as nM concentration) were calculated by plotting the inhibition rate against the antibody concentration in GraphPad Prism V8.0 software. UR, scFv unrelated; CS, Covid Serum pool; *Data expressed as dilution giving 50% inhibition.

**Table S7. Secondary structure content estimation of scFv76-cluster antibodies derived from the circular dichroism spectra at 20 °C**

| scFv | α-helix (%) | Antiparallel β-sheet (%) | Parallel β-sheet (%) | β-turn (%) | Other (%) |
|---|---|---|---|---|---|
| **76** | 1.1 | 39.0 | 0.0 | 14.7 | 45.2 |
| **76-46** | 0.0 | 45.7 | 0.0 | 13.6 | 40.7 |
| **76-55** | 0.0 | 43.1 | 0.0 | 13.1 | 43.8 |
| **76-57** | 0.0 | 41.7 | 0.0 | 12.8 | 45.4 |
| **76-58** | 0.1 | 43.8 | 0.0 | 12.5 | 43.6 |
| **76-77** | 1.5 | 42.8 | 0.0 | 12.4 | 43.3 |

**Figure S1**

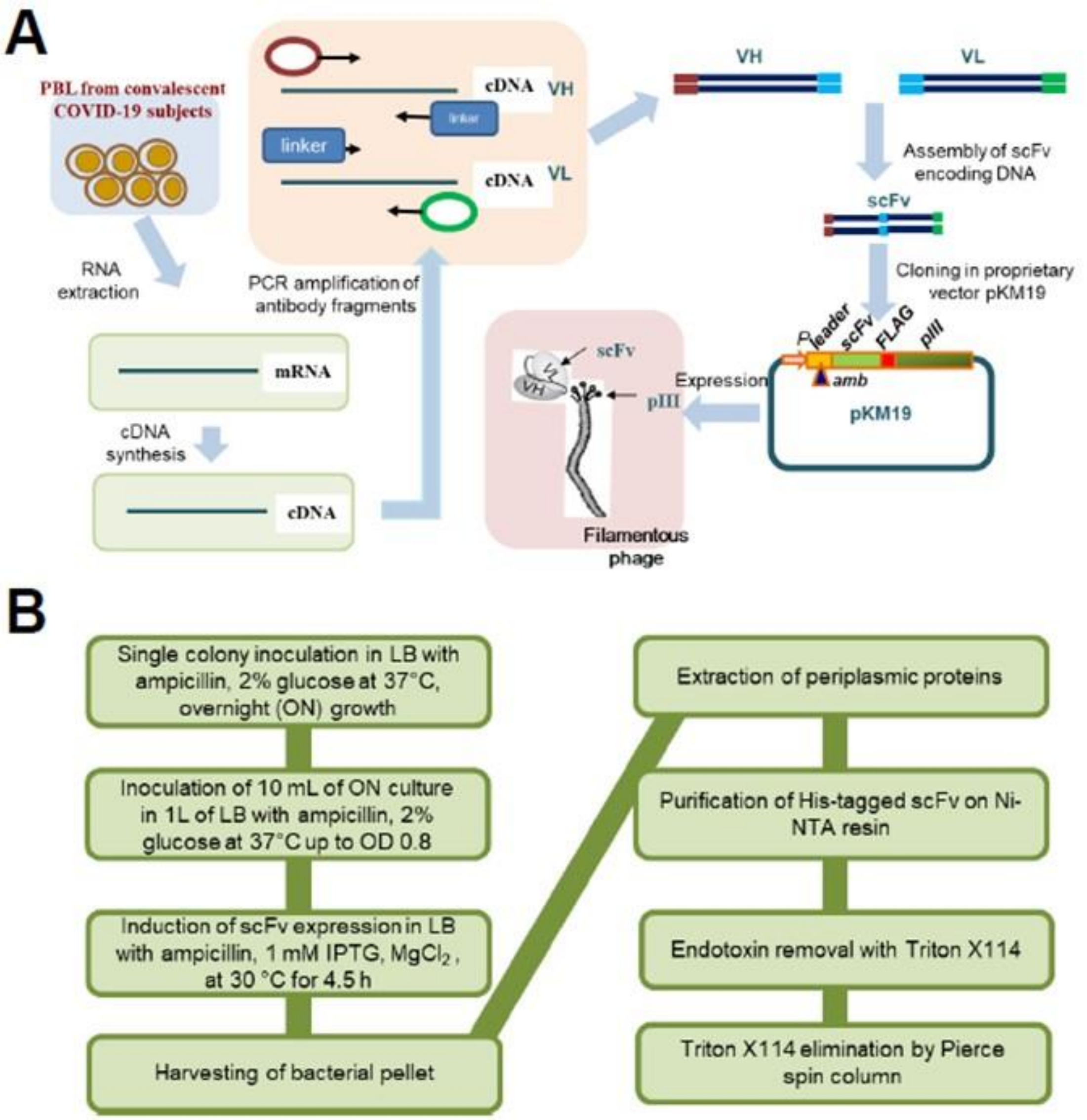


**Schematic representation of experimental procedure to select anti-SARS-CoV-2 spike RBD human scFv antibodies.** (**A**) Construction of phage display libraries. (**B)** Flow chart of production method of soluble scFv antibodies.

**Figure S2**

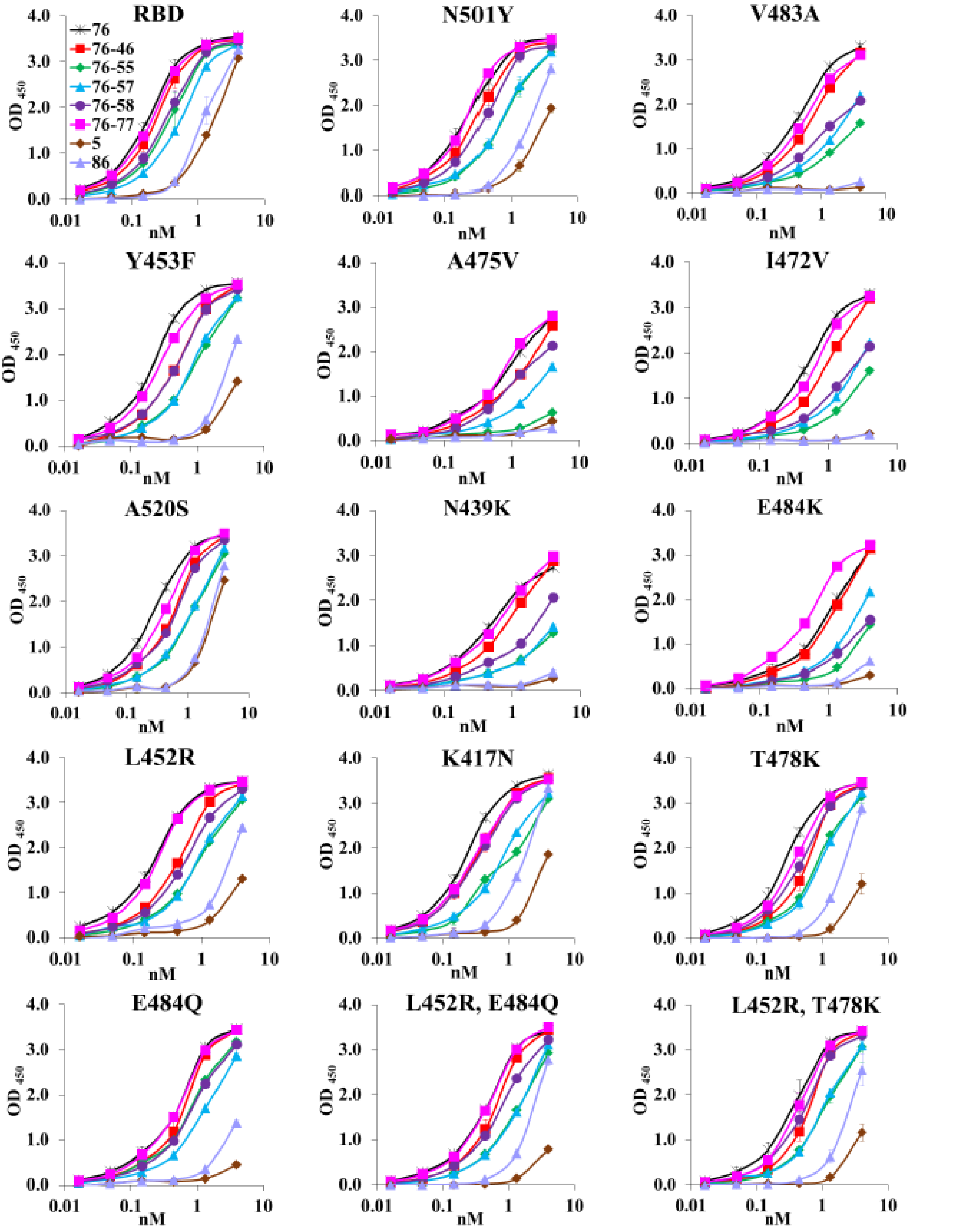


**Evaluation of the effect of RBD mutations on scFv antibody binding**. Binding of scFv antibodies to mutated RBD proteins by ELISA. RBD: Wuhan strain. Data are the average (± SD) of three technical replicates from one representative experiment.

**Figure S3**

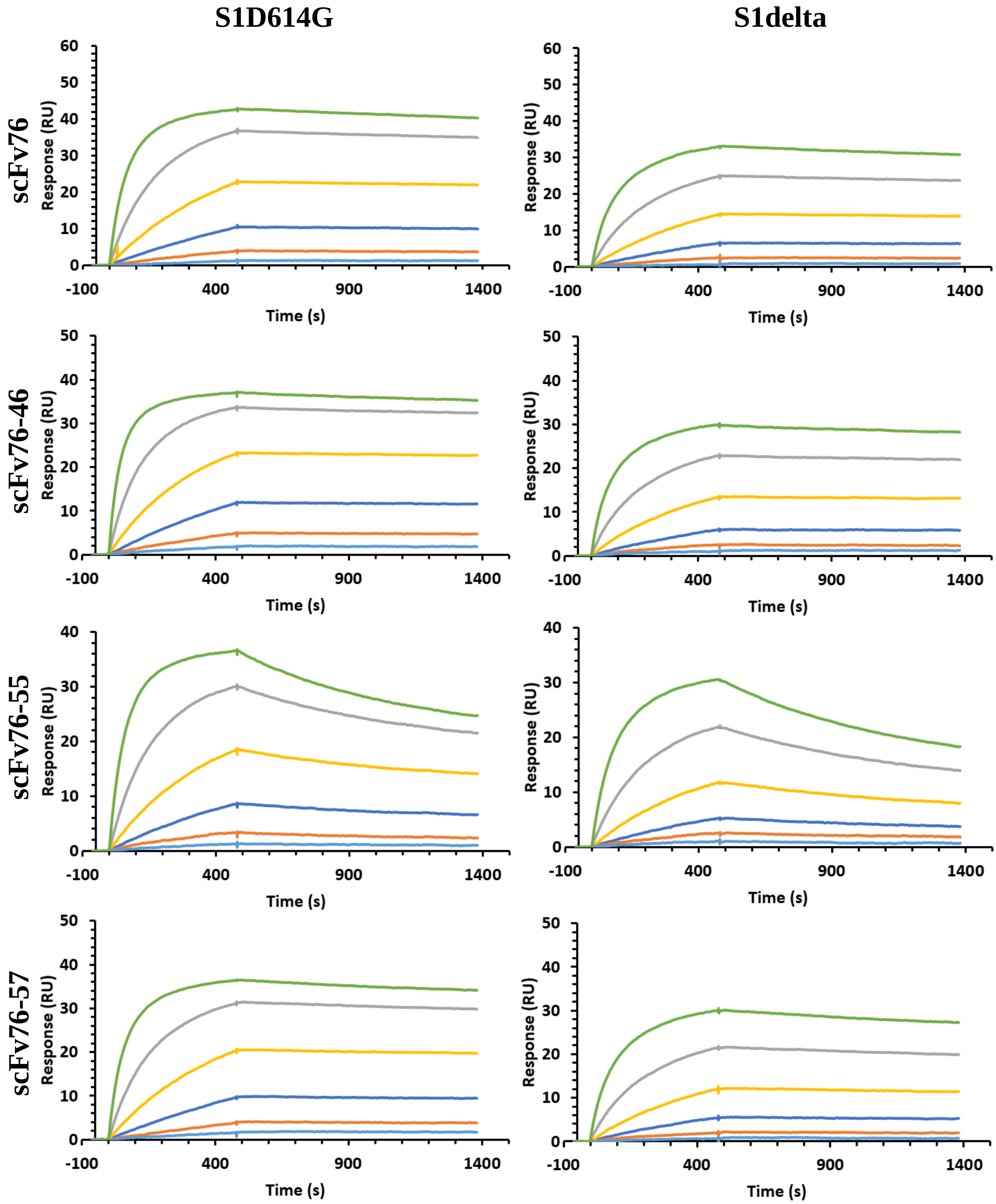


Continues…

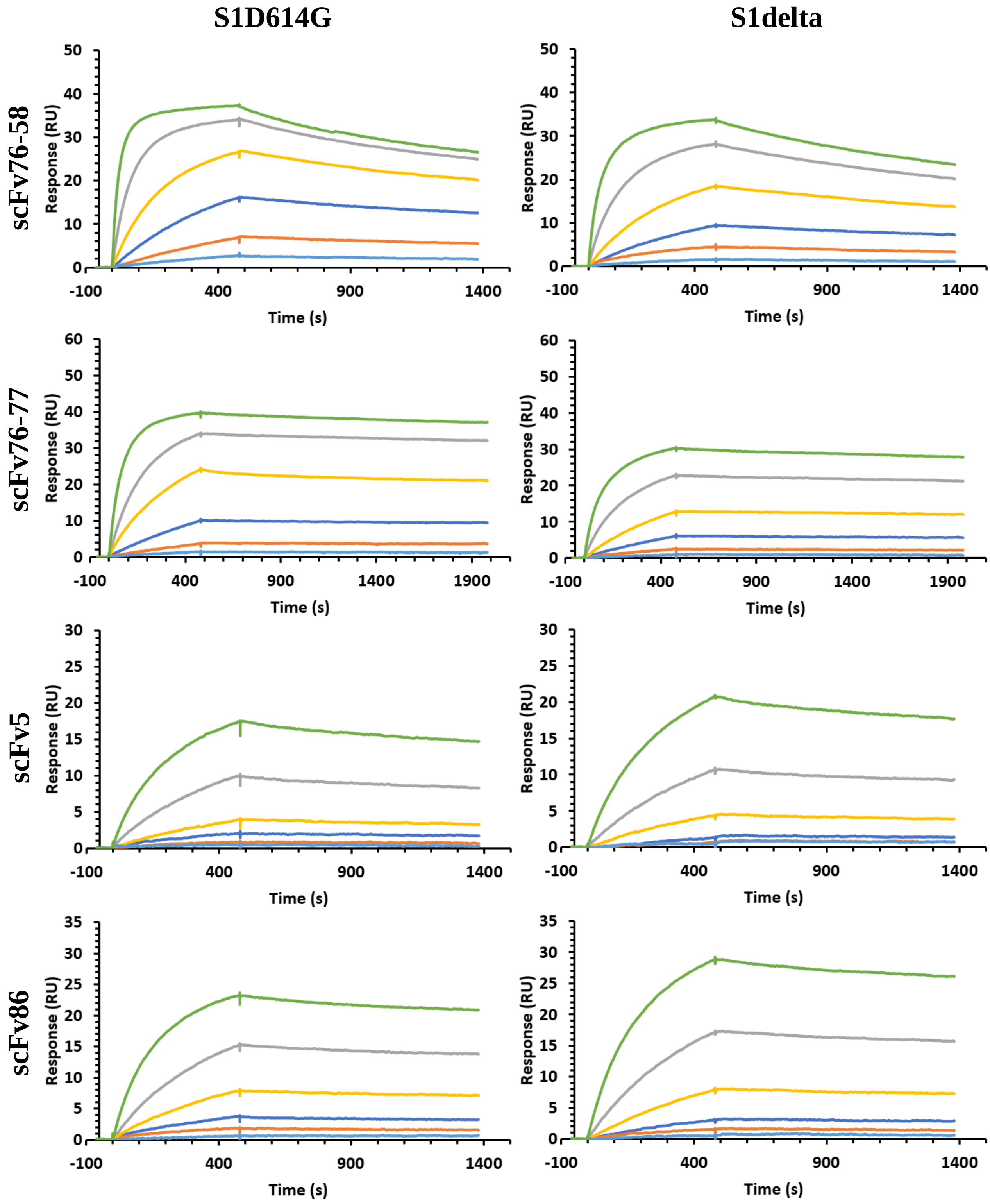


**SPR analysis of interaction of indicated antibodies with spike Subunit 1 proteins**. Left column S1 with mutation D614G (S1D614G), right column S1 with mutations T19R, G142D, E156G, 157-158d, L452R, T478K, D614G, P681R, as in Delta (B.1.617.2) VOC (S1delta). Concentrations: green 113 nM, gray 37.67 nM, yellow 12.56 nM, blue 4.19 nM, orange 1.40 nM and light blue 0.47 nM.

**Figure S4**

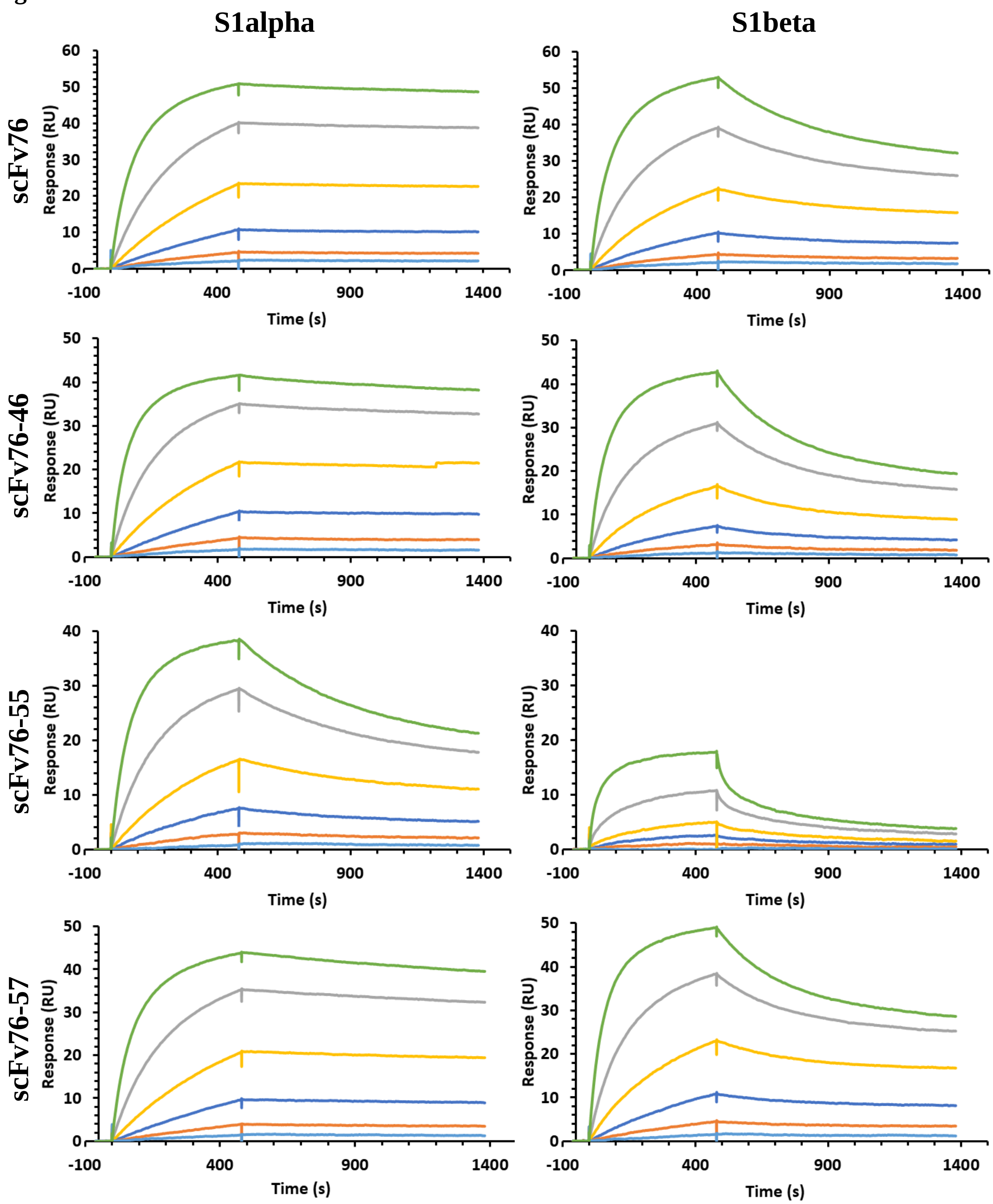


Continues…

Continues Figure S4

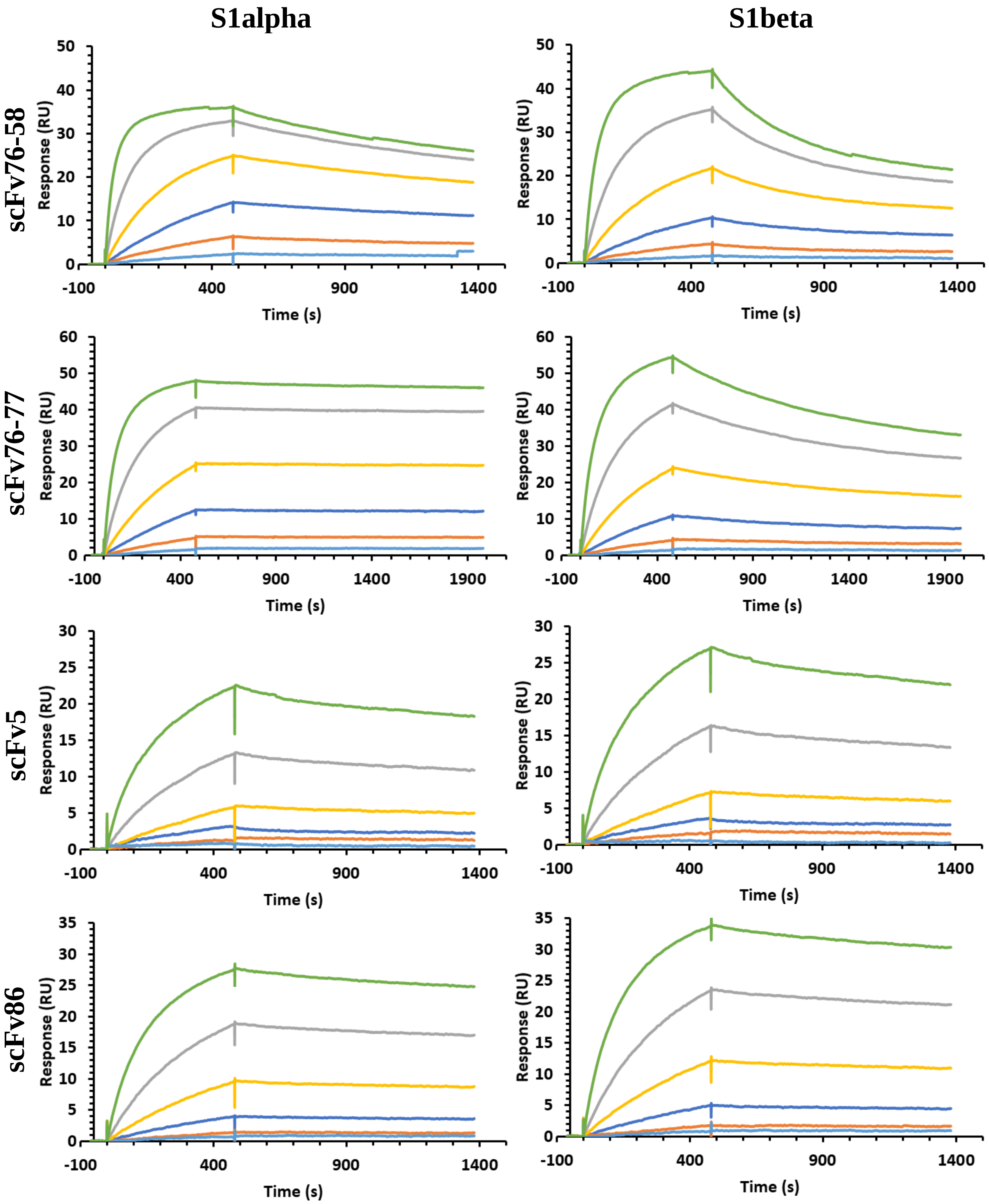


**SPR analysis of interaction of indicated antibodies with spike Subunit 1 proteins.** Left column S1 with mutations HV69-70d, Y144d, N501Y, A570D, D614G, P681H, as in Alpha (B.1.1.7) VBM (S1alpha); right column S1 with mutations K417N, E484K, N501Y, D614G, as in Beta (B.1.351) VBM (S1beta). Concentrations: green 113 nM, gray 37.67 nM, yellow 12.56 nM, blue 4.19 nM, orange 1.40 nM and light blue 0.47 nM.

**Figure S5**

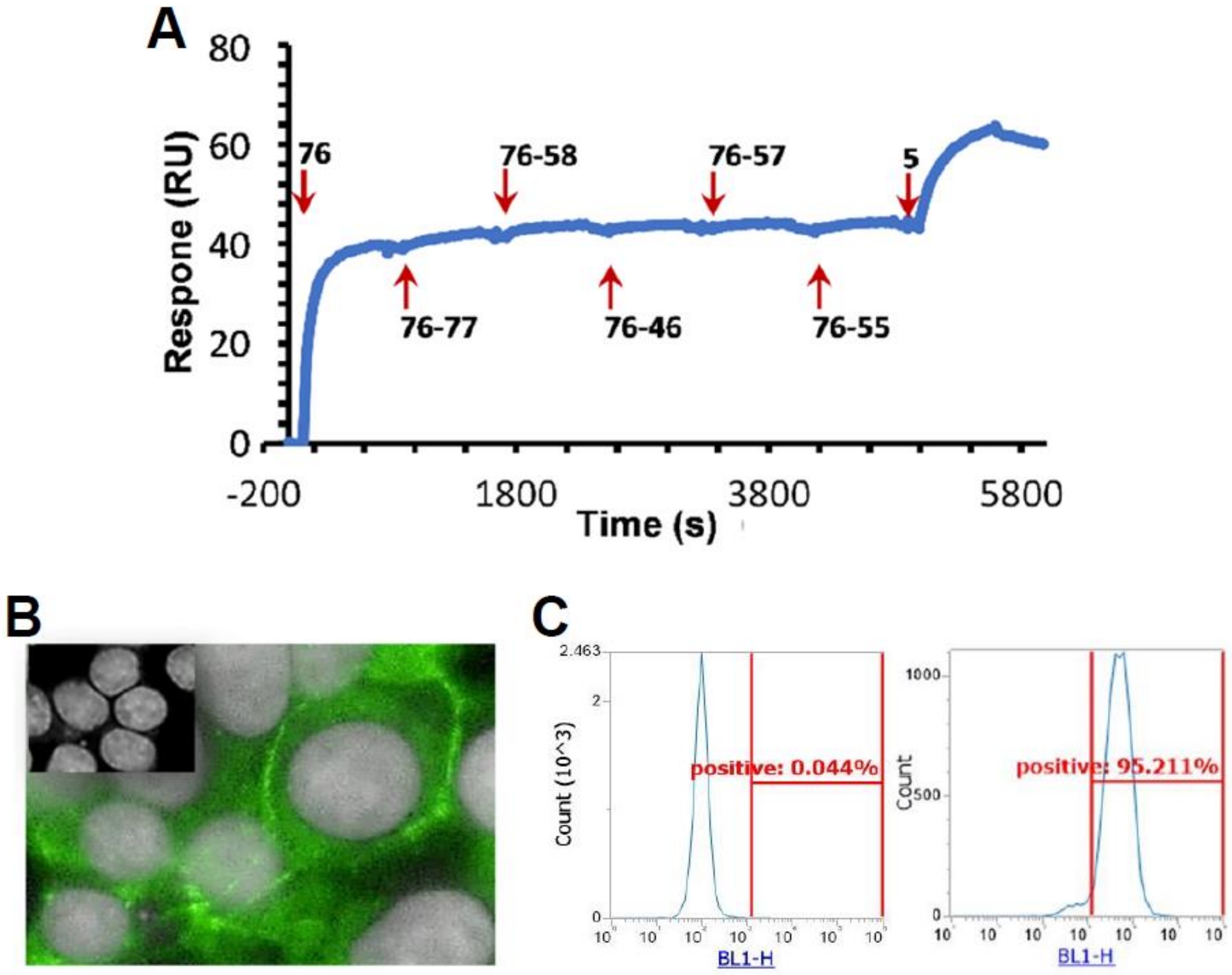


**Characterization of 76clAbs binding. (A)** SPR epitope binning by injection of scFv76 antibody followed by indicated scFv antibodies. (**B)** High content screening immunofluorescence imaging (Operetta) of spike-transfected HEK293T cells incubated with scFv76 antibody. Binding detected by FITC-conjugated anti-tag antibody. Inset, mock transfected cells. (**C**) FACS analysis of mock-transfected (left) or spike-transfected (right) HEK293T cells, following treatment with 360 nM scFv76 antibody, for 1h at 4°C. Detection through mouse anti-Flag M2-FITC.

**Figure S6**

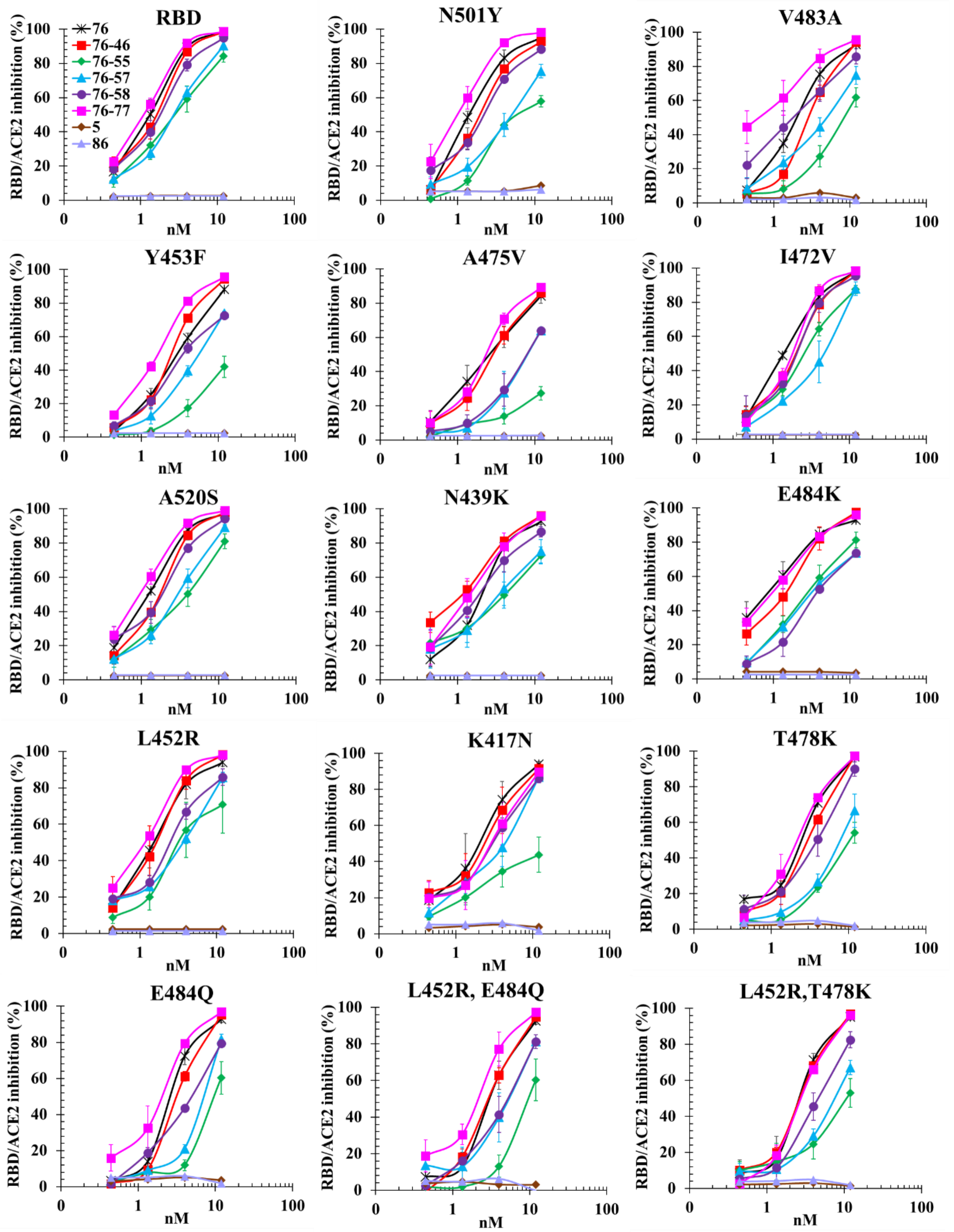


**Evaluation of the effect of RBD mutations on scFv antibody inhibitory activity of RBD/ACE2 interaction.** Inhibition of spike/ACE2 interaction by scFv antibodies measured by ELISA. RBD: Wuhan strain. Data are expressed as percentage inhibition and are the average (± SE) of 2-3 independent experiments.

**Figure S7**

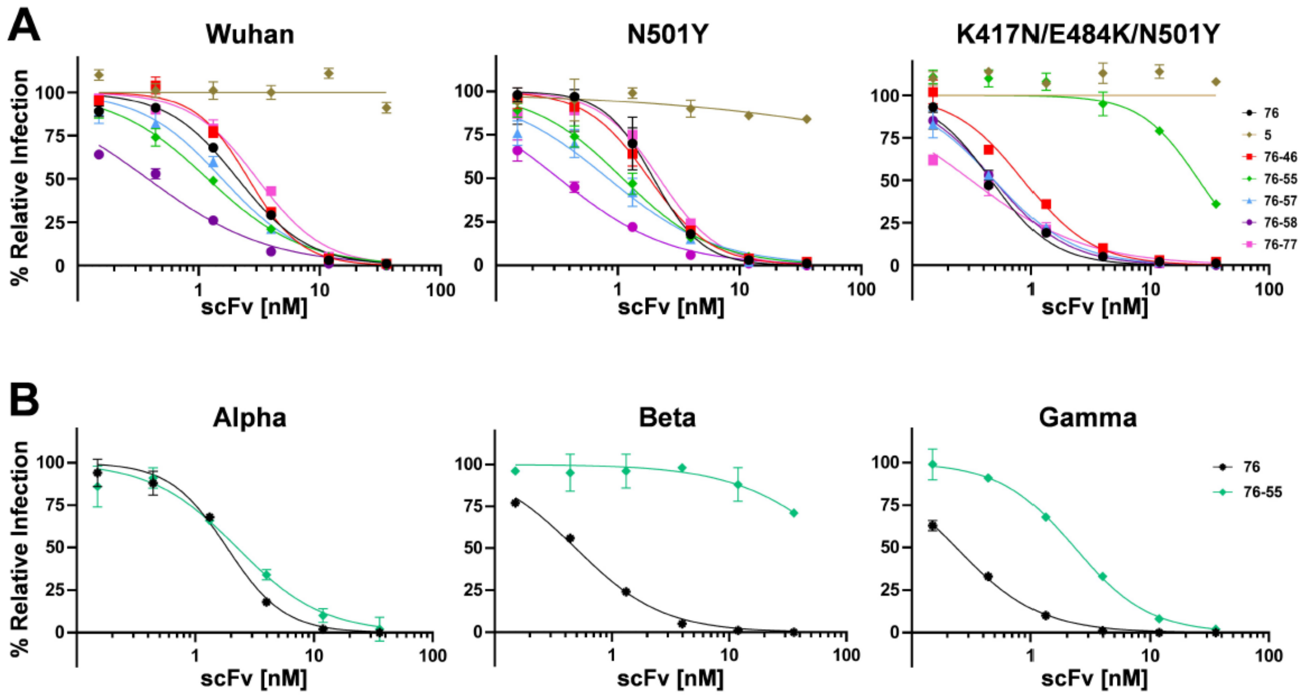


**Neutralization of pseudotyped-virus expressing indicated spikes assessed by luciferase-assay in hACE2-expressing Caco-2 cells.** Data are the average (±SD) of two replicates from one representative experiment.

### Figure S8

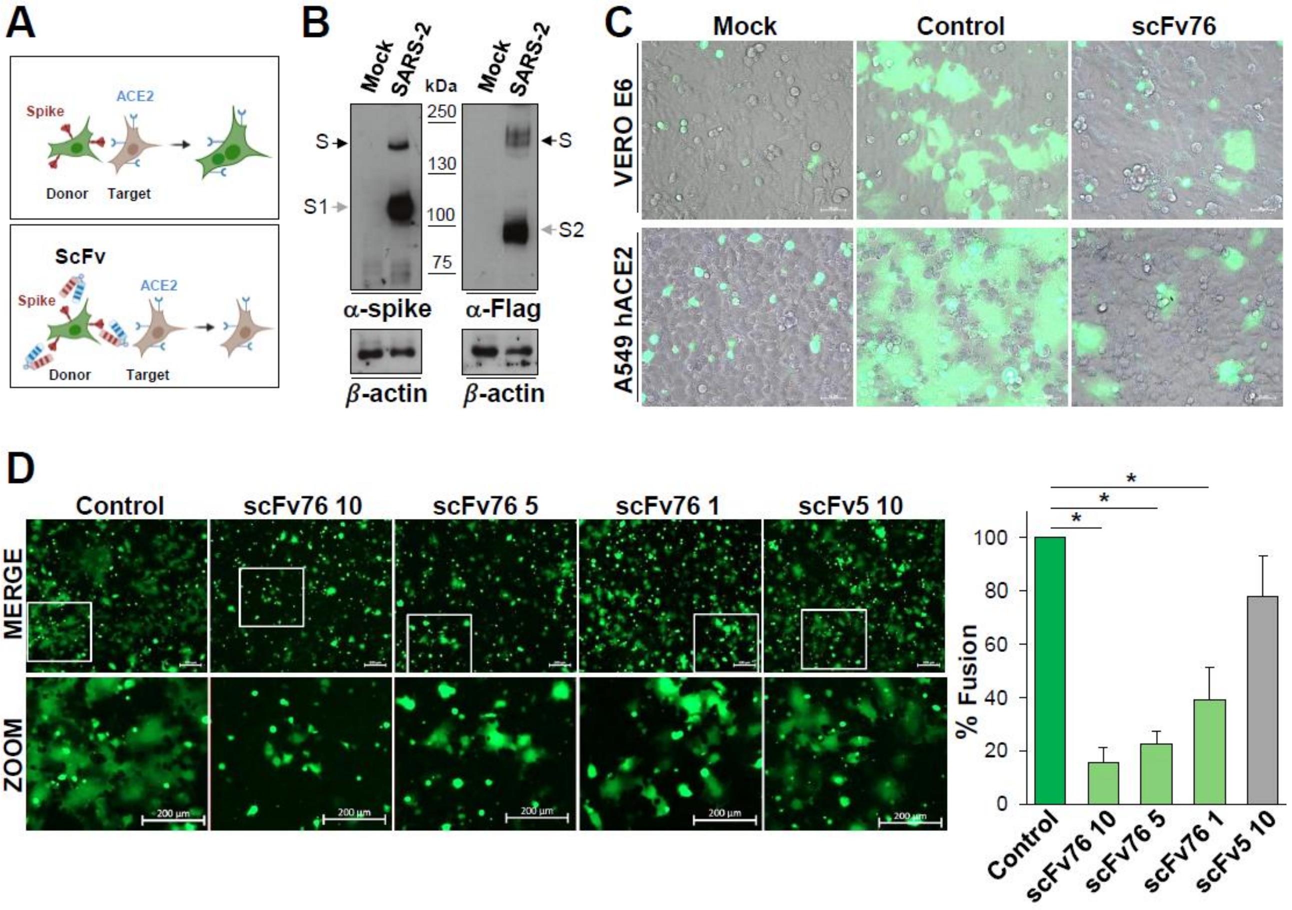


**Inhibition of SARS-CoV-2 S-mediated cell-cell fusion by scFv76 antibody**. HEK293T cells (donor cells) were co-transfected with plasmids encoding Flag-tagged SARS-CoV-2 spike (SARS-2, Wuhan) and GFP, or empty vector and GFP (Mock). (**A**) Schematic representation of scFv-mediated inhibition of syncytia formation in the donor-target cell-cell fusion assay. (**B)** Spike protein levels in donor cells detected by immunoblot using anti-spike or anti-Flag antibodies as described (*29*) are shown. Black arrows indicate bands corresponding to uncleaved S proteins, whereas gray arrows indicate bands corresponding to the S1 or S2 subunits. (**C**) Donor cells were detached 40 h post transfection, incubated with scFv76 (10 μg/mL) for 30 min, and overlaid on Vero E6 (top) or A549-hACE2 (bottom) cell monolayers. After 4 h, cell-cell fusion was assessed by fluorescence microscopy. Merge images are shown. Scale bar 50 μm. (**D)** Donor cells incubated with scFv76 (10, 5 or 1 μg/mL) or scFv5 (10 μg/mL) antibodies for 30 min were overlaid on A549-hACE2 cell monolayers. After 4 h, cell-cell fusion was assessed as in **C**. Fluorescence images are shown. Scale bar, 200 μm (zoom, 200 μm). Cell-cell fusion was determined and expressed as percentage relative to control (right panel). Data represent the average (± SD) of 5 fields from two biological replicates. *P<0.0001; ANOVA test.

**Figure S9**

A

| | VOC | VBMs | | | | | | | | |
|---|---|---|---|---|---|---|---|---|---|---|
| | Delta B1.617.2 | Alpha B.1.1.7 | Beta B.1.351 | Gamma P.1 | Epsilon B.1.427/9 | Eta B1.525 | Iota B1.526 | Kappa B1.617.1/3 | Zeta P.2 | Mu B.1621 |
| R346 | | | | | | | | | | K |
| K417 | N* | | N | T | | | | | | |
| L452 | R | | | | R | | R* | R | | |
| S477 | | | | | | | N* | | | |
| T478 | K | | | | | | | | | |
| E484 | | K* | K | K | | K | K | Q | K | K |
| S494 | | P* | | | | | | | | |
| N501 | | Y | Y | Y | | | | | | Y |
| D614 | G | G | G | G | G | G | G | G | G | G |

*Found in some isolates

B

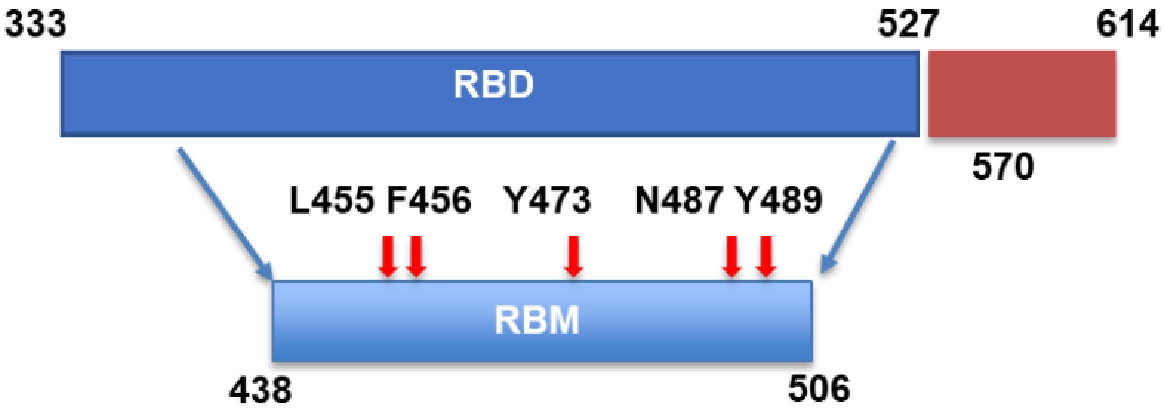


**SARS-CoV-2 spike RBD and RBM. (A)** Mutations found in Variant of Concern (VOC) and Variants Being Monitored (VBMs). Adapted from CDC SARS-CoV-2 Variant Classifications and Definitions (cdc.gov) update October 4th, 2021. **(B)** Schematic representation of spike RBD and RBM. Red arrows indicate the amino acid residues of the antigenic epitope recognized by 76clAbs none of which is found in SARS-CoV-2 variants.

**Figure S10**

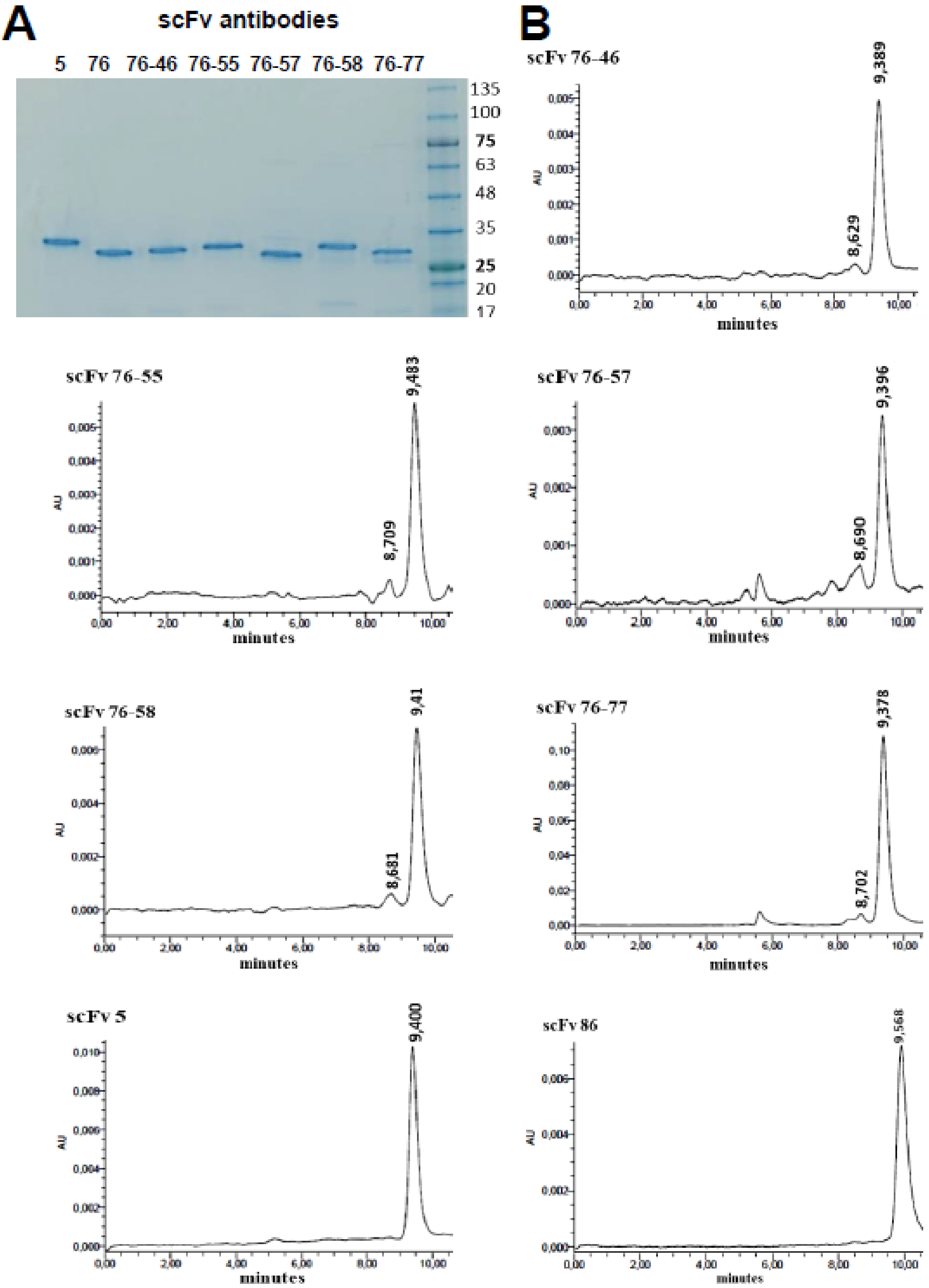


**Biochemical characterization of scFv76-cluster antibodies. (A)** SDS-PAGE analysis (Last lane molecular weight markers (kDa). **(B)** UV chromatogram at 280 nm from SEC-HPLC analysis of scFv76 in 50 mM phosphate buffer, 150 mM NaCl, 10% acetonitrile, pH 7.2 on BioSep SEC-s2000 (Phenomenex) column.

**Figure S11**

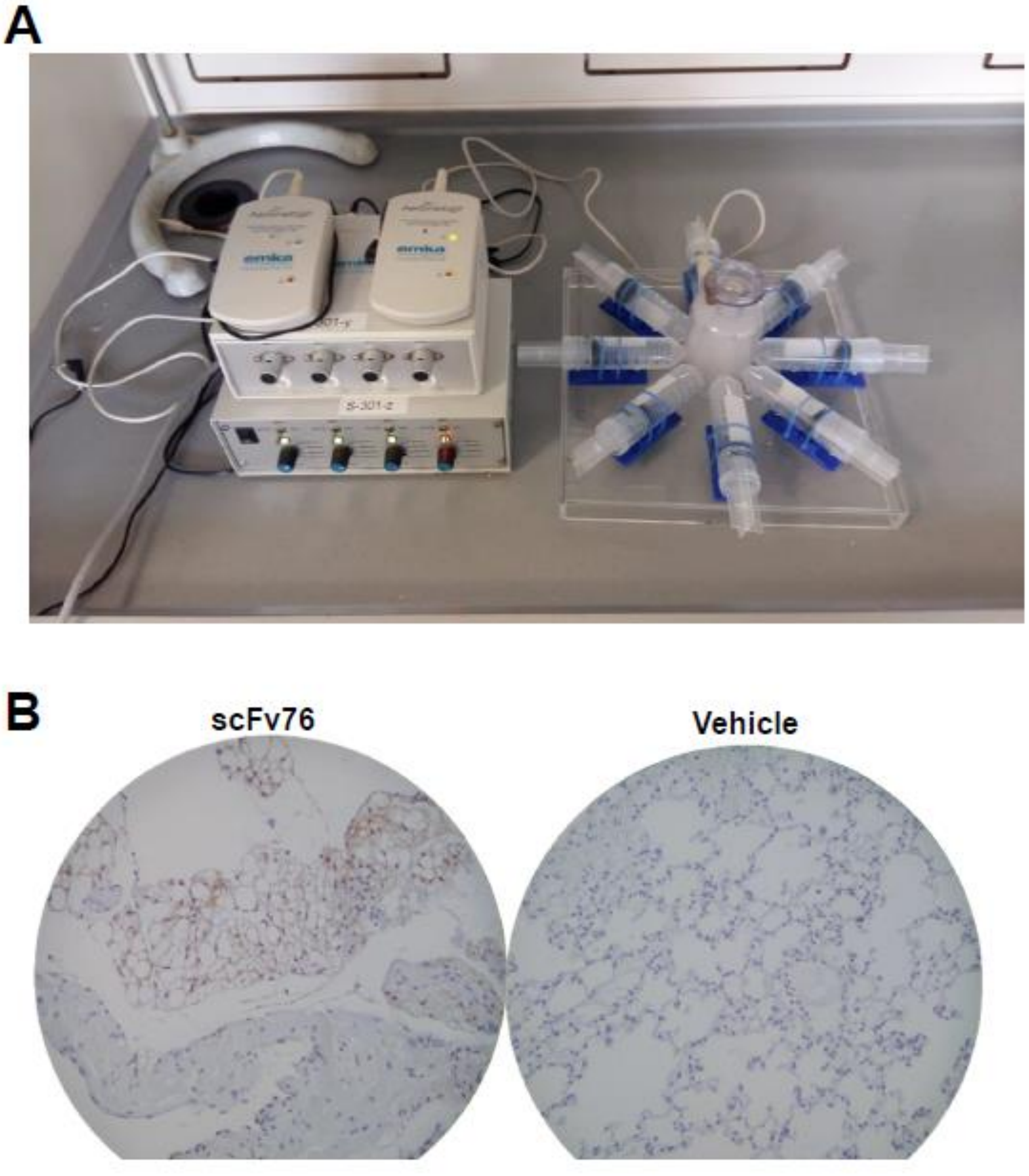


**Equipment for nebulization of 76clAbs and nose-only delivery to mice**. **(A)** Equipment was built by adapting the nebulization device Aerogen Pro mesh nebulizer, approved for human aerosol treatment, to a nose-only inhalation chamber for mice. **(B)** Pulmonary localization of scFv76 in mice 1h after nebulization of 5 mg/mL solution assessed by immunohistochemistry. Antibody accumulations are stained in brown.

**Figure S12**

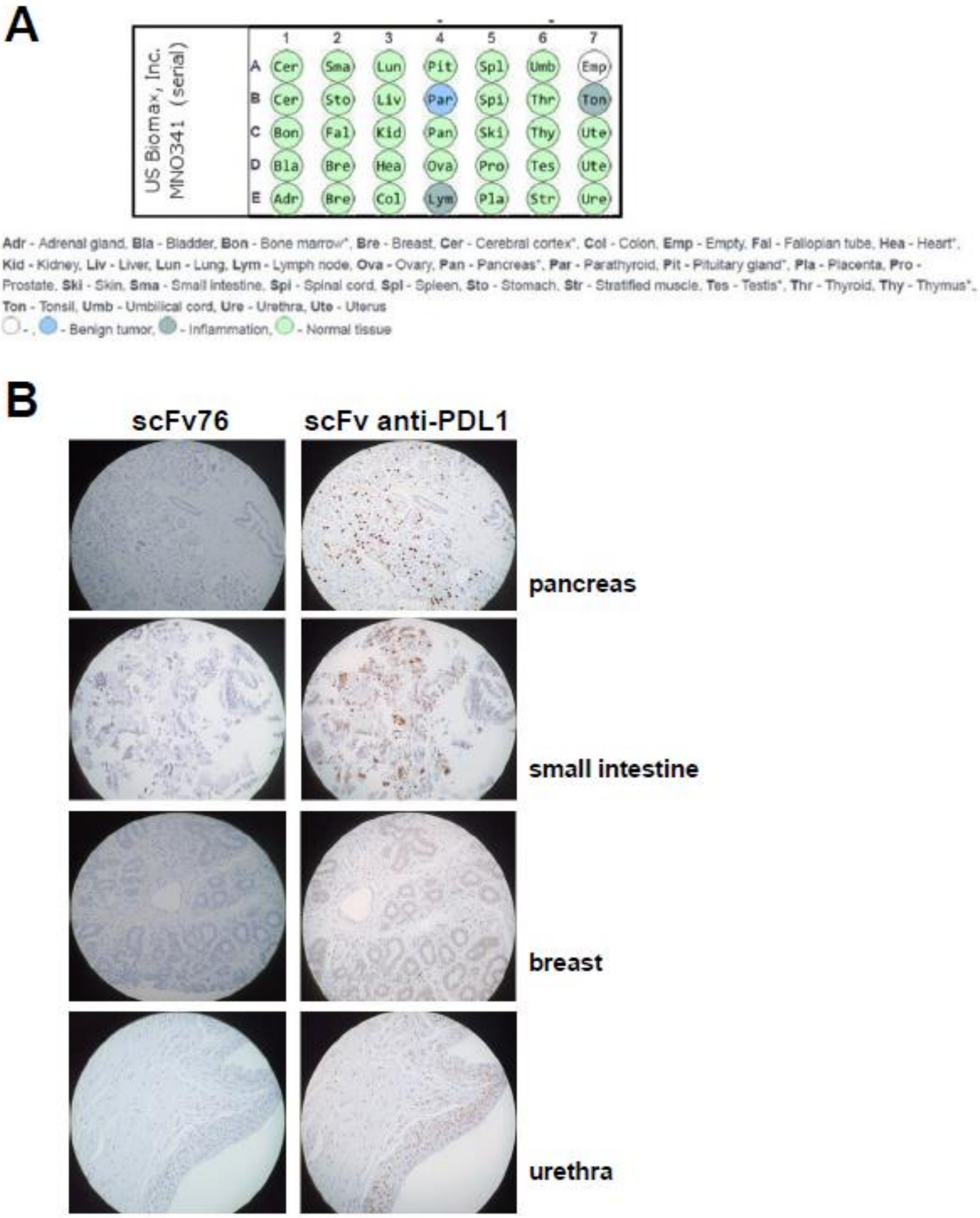


**scFv76 does not react with human normal tissues.** (**A**) Tissue microarray. (**B**) Representative immunohistochemistry images. An anti-hPDL1 scFv was used as positive control. Detection by anti-His-Tag antibody.
